# Fibulin-2 transduces a matrix-to-metabolism signal in kidney fibrosis

**DOI:** 10.64898/2026.06.19.733428

**Authors:** Yuan Gui, Yuanyuan Wang, Wenxue Li, Jia-Jun Liu, Chen Dai, Samantha Mae Mallari, Kelly Zheng, Cameron Jones, Henry Wells Shaffer, Liora Yueran Dorsett, Tessora Chang, Benjamin Malowitz, Yanbao Yu, Wei Chen, Silvia Liu, Hongbo Liu, Yansheng Liu, Dong Zhou

**Affiliations:** Division of Nephrology, Department of Medicine, University of Connecticut School of Medicine, Farmington, CT, 06030, USA; Yale Cancer Biology Institute, Yale University, West Haven, Connecticut, 06520, USA; Department of Pharmacology, School of Medicine, Yale University, New Haven, Connecticut, 06520, USA; Organ Pathobiology and Therapeutics Institute, University of Pittsburgh School of Medicine, Pittsburgh, PA, 15261, USA; Department of Chemistry & Biochemistry, University of Delaware, Newark, DE, 19716, USA; Nephrology Division, Department of Medicine, Albert Einstein College of Medicine, Bronx, NY, 10461, USA; Department of Pharmacology and Chemical Biology, University of Pittsburgh School of Medicine, Pittsburgh, PA, 15261, USA; Department of Biomedical Genetics, University of Rochester Medical Center, Rochester, NY, 14642, USA; Department of Biomedical Informatics and Data Science, Yale University School of Medicine, New Haven, Connecticut. 06516, USA

**Keywords:** Smoothened, Fibulin 2, ACAT1, GWAS, Proteomics, Lipidomics

## Abstract

Fibrotic extracellular matrix (ECM) is not merely a structural scaffold but an instructive signaling interface that shapes epithelial cell state. However, the molecular cues by which matrix remodeling controls tubular metabolism during kidney fibrosis remain poorly defined. Here, we identify Fibulin-2 (FBLN2) as a fibroblast-derived matrix cue that transduces fibrotic ECM remodeling into tubular mitochondrial metabolic reprogramming. Using fibroblast-selective deletion of Smoothened (Smo) across distinct fibroblast subpopulations, we found that loss of fibroblast Smo preserved kidney function and attenuated fibrosis in mouse models of chronic kidney injury. Multi-omics profiling revealed coordinated remodeling of the fibrotic matrisome, highlighted by suppression of FBLN2, an ECM glycoprotein genetically linked to kidney function in humans. Mechanistically, FBLN2 engaged EGFR in tubular epithelial cells and activated EGFR-AKT signaling in a non-canonical ligand-like manner. This signaling axis suppressed acetyl-CoA acetyltransferase 1 (ACAT1), a mitochondrial regulator of fatty acid oxidation and amino acid metabolism. Disruption of fibroblast Smo-FBLN2 signaling restored ACAT1-dependent oxidative metabolism and reduced tubular fibrotic activation. Spatial lipidomics revealed compartment-specific lipid remodeling associated with altered mitochondrial fatty acid metabolism, including acylcarnitine and phospholipid changes linked to reduced fibrotic injury. Together, these findings define a Fibulin-2-EGFR-ACAT1 matrix-to-metabolism signaling axis that couples fibrotic ECM remodeling to tubular mitochondrial metabolism during kidney fibrosis.

## Introduction

Beneath the visible scars of chronic disease lies a hidden code: a dynamic interplay of extracellular matrix (ECM), metabolism, and signaling that determines whether tissues heal or collapse(*1, 2*). Fibrosis, long dismissed as a passive ECM accumulation, is now recognized as a metabolically active process that relentlessly damages organs, including the kidney, liver, lung, heart, and brain, imposing a staggering global disease burden(*3, 4*). In the kidney alone, chronic kidney disease (CKD) is projected to rank among the world’s top five causes of death by 2040(*5*), highlighting fibrosis remains a pressing and unsolved medical challenge.

The kidney’s extraordinary metabolic demand, second only to the heart(*6*), is fueled predominantly by proximal tubular epithelial cells (PTECs), which reclaim most of the glomerular filtrate. To sustain this workload, PTECs rely on mitochondrial fatty acid oxidation (FAO) to generate ATP through oxidative phosphorylation (OXPHOS), making lipids their principal energy source(*7, 8*). This specialization renders proximal tubules highly vulnerable to hypoxia and energetic stress. Injury impairs mitochondrial function, suppressing FAO and ketogenesis(*9–12*), reducing lipid utilization, and driving intracellular lipid accumulation(*13*). Glycolysis and alternative substrates, such as glutamine, can transiently compensate(*6*), but cannot match the efficiency of lipid-derived ATP. Heightened glycolytic flux generates cytosolic acetyl-CoA and NADPH, feeding de novo lipogenesis(*14*), and aggravating lipotoxic stress. Parallel pathways, including amino acid metabolism and gluconeogenesis(*15, 16*), intersect with lipid oxidation to maintain energy equilibrium. Persistent disruption of these intertwined metabolic programs triggers lipotoxicity, inflammation, tubular dysfunction, and maladaptive repair, accelerating progression to fibrosis(*15, 17*).

While tubular injury sets the stage for CKD(*18*), the fibrotic ECM directs its trajectory(*19*). ECM proteins, largely secreted by fibroblasts, actively influence mitochondrial function and OXPHOS efficiency(*10, 20, 21*), suggesting that the matrix itself encodes a metabolic language. Our prior work demonstrated that fibroblast-expressed Smoothened (SMO), an atypical G-protein-coupled receptor (GPCR) at the core of Hedgehog signaling, remodels the matrix after acute kidney injury (AKI)(*22*). During this process, proteomics revealed that differentially expressed proteins are mainly involved in mechanical stress and defense response at early phase, whereas metabolic changes, particularly mitochondrial FAO, emerge during tubular repair and regeneration and become prominent as disease progresses toward CKD(*8, 13, 15*). However, how fibroblasts regulate tubular mitochondrial FAO in a fibrotic microenvironment and which matrix-derived factors impact metabolism during disease transition, remain unclear. Here, we investigated whether fibroblast-intrinsic SMO acts as a molecular hub linking ECM remodeling to tubular lipid metabolism, thereby directing maladaptive repair. Deciphering this Smo-ECM-lipid axis promises to reveal how fibroblasts coordinate structural and metabolic programs to drive fibrotic transition.

## Results

### Fibulin-2 mediates anti-fibrotic matrix remodeling downstream of Gli1+ fibroblast *Smo*

To define the role of fibroblast-SMO in kidney fibrosis, we crossed *Smo*-floxed mice with *Gli1-CreER^T2^* mice to generate conditional knockout mice lacking *Smo* in Gli1+ fibroblasts (*Gli1*-*Smo*-/-). Littermate *Smo*-floxed mice lacking Cre (*Gli1*-*Smo*+/+) served as controls. Tamoxifen was administered for five consecutive days to induced recombination(*22*), followed by a one-week washout before induction of fibrosis using either unilateral ischemia-reperfusion injury with contralateral nephrectomy (UIRI + UNx, 17 days) or unilateral ureter obstruction (UUO, 7 days) (Figure 1A).

**Figure 1:**
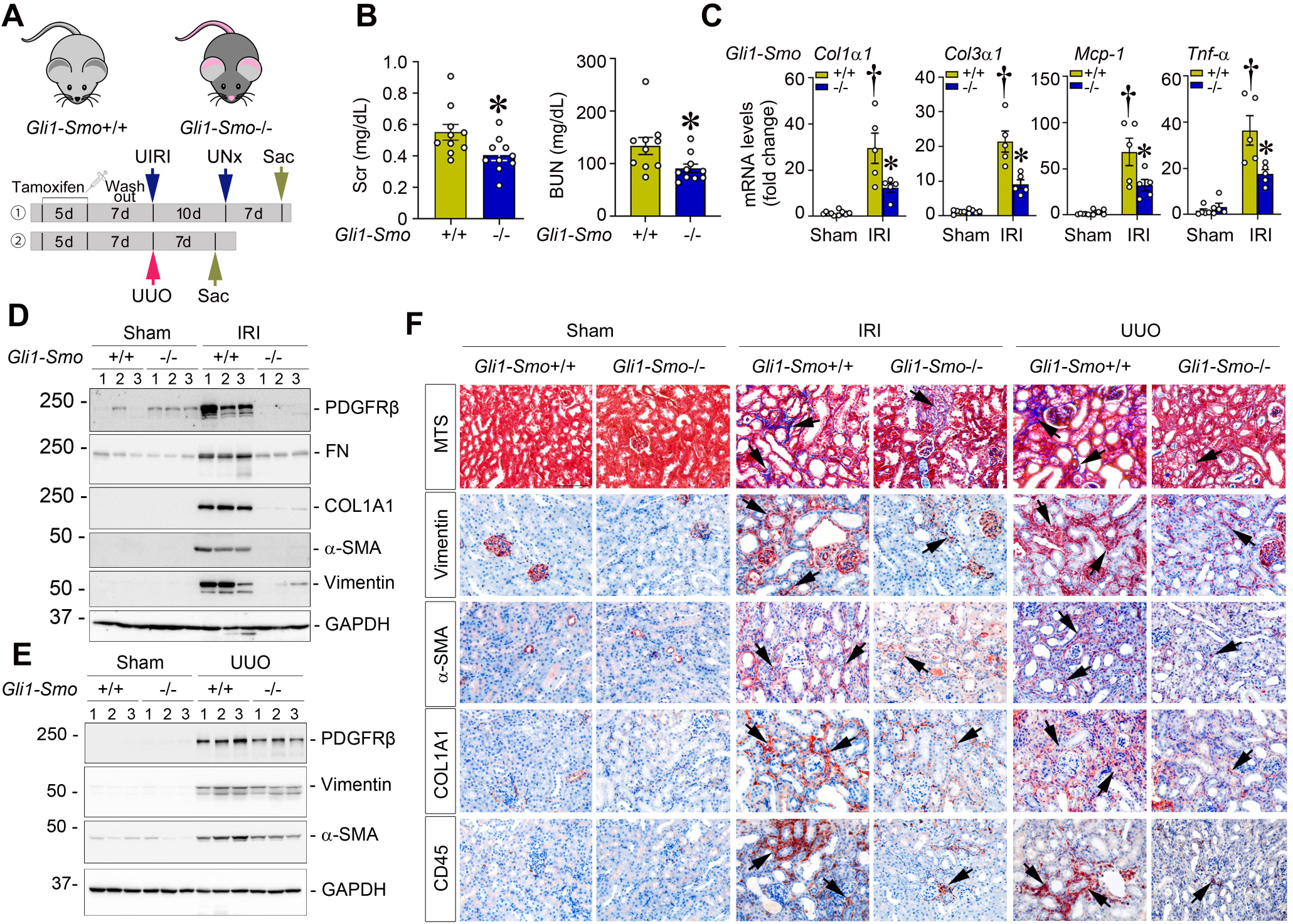
Fibroblast-specific ablation of *Smo* ameliorates kidney fibrosis. (**A**) Experiment design. *Gli1-Smo*+/+ and *Gli1-Smo*-/- mice were administered tamoxifen prior to unilateral ischemia-reperfusion injury (UIRI) combined with nephrectomy (model ①) or unilateral ureter obstruction (UUO, model ②). In model ①, the contralateral kidney was removed 10 days (UNx) after UIRI, and mice were sacrificed 7 days thereafter. In model ②, mice were sacrificed 7 days after UUO. After UIRI combined with UNx, (**B**) Serum creatinine (Scr) and Blood Urea Nitrogen (BUN) levels (n = 10), (**C**) qPCR of *Col1a1*, *Col3a1*, *Mcp-1*, and *Tnf-a* mRNA (n = 5), (**D**) Western blotting showing platelet-derived growth factor receptor β (PDGFRβ), fibronectin (FN), COL1A1, α-SMA, and Vimentin proteins in *Gli1-Smo*+/+ and *Gli1-Smo*-/- kidneys after UIRI + UNx. (**E**) Western blotting showing PDGFRβ, Vimentin, and α-SMA proteins in *Gli1-Smo*+/+ and *Gli1-Smo*-/- mice after UUO. (**F**) Representative micrographs of Masson’s Trichrome staining (MTS) and immunohistochemical staining for Vimentin, α-SMA, COL1A1, and CD45 in *Gli1-Smo*+/+ and *Gli1-Smo*-/- kidneys after UIRI + UNx and UUO. Scale bar, 25 µm. Arrows indicate positive cells. † *P* < 0.05 versus sham control, * *P* < 0.05 versus *Gli1-Smo*+/+ UIRI+UNx mice. Graphs are presented as means ± SEM. Differences among groups were analyzed using two-sided unpaired t-tests or one-way ANOVA followed by the Student-Newman-Keuls test.

In the ischemic model, *Gli1-Smo*-/- mice exhibited preserved kidney function with significant lower serum creatinine (Scr) and blood urea nitrogen (BUN) levels compared with controls (Figure 1B). Fibrosis and inflammation markers, including α1 type I collagen (*Col1α1)*, α3 type III collagen (*Col3α1*), monocyte chemoattractant protein-1 (*Mcp-1)*, and tumor necrosis factor-α (*Tnf-α*), were substantially reduced (Figure 1C). Western blotting confirmed decreased platelet-derived growth factor receptor β (PDGFRβ), fibronectin (FN), COL1Al, α-Smooth Muscle Actin (α-SMA), and Vimentin in both models (Figure 1, D and E; Supplemental Figure 1, A and B). Immunostaining and Masson’s trichrome staining (MTS) further demonstrated reduced collagen deposition, myofibroblasts activation (α-SMA+, Vimentin+), and monocytes infiltration (CD45+) in *Gli1-Smo*-/- kidneys (Figure 1F; quantification in Supplemental Figure 1, C-G). Together, these results show that *Smo* deletion in Gli1+ fibroblasts attenuates kidney fibrosis in both ischemic and obstructive injury models.

To uncover the molecular basis, we performed data-independent acquisition (DIA)-based proteomics on ischemic kidneys. Principal component analysis (PCA) revealed clear segregation between *Gli1-Smo*+/+ and *Gli1-Smo*-/- kidneys (Figure 2A). Among 8,024 identified proteins, 5,945 were differentially expressed using the P value cutoff of 0.05 (Figure 2B; replicate correlations and the intensity distributions shown in Supplemental Figure 1, H and I). Gene ontology (GO) analysis highlighted ECM organization and assembly as the most significant altered biological processes in *Gli1-Smo*-/- kidneys (Figure 2C). Matrisome analysis identified 51 differentially expressed secreted core matrisome proteins (Figure 2D), including structural collagens (COL1A1, COL1A2, COL3A1, COL12A1, etc), basement membrane components (LAMA/LAMB/LAMC1 family, NID1, NID2, etc), and matricellular regulators such as fibulins (FBLN), tenascin-C (TNC), and periostin (POSTN). Among them, fibulins emerged as prominent candidates. Fibulins are ECM glycoproteins that organize collagen, FN, and basement membrane components to stabilize matrix architecture during tissue remodeling(*23*). Volcano plot analysis revealed significant downregulation of FBLN1, FBLN2, and FBLN7 in *Gli1-Smo*-/- kidneys (Figure 2E), with FBLN2 showing the most significant reduction (Figure 2, F and G), confirmed by quantitative PCR (qPCR) and western blotting (Figure 2, H and I; Supplemental Figure 2A). Reanalyzing the public single-nucleus RNA sequencing dataset (GSE139107)(*24*), we showed that fibroblasts and endothelial cells are the principal *Fbln2*-expressing cells in mouse fibrotic kidneys (Figure 2J). Consistently, we observed elevated *FBLN2* expression in fibroblasts and endothelial cells from human CKD kidneys, using the Kidney Precision Medicine Project (KPMP) dataset(*25*) (Supplemental Figure 2, B and C). Chromatin accessibility analysis revealed increased open chromatin at the *Fbln2* promoter in fibroblasts and endothelial cells in mouse kidneys(*26*) (Supplemental Figure 2D). Importantly, using the genome-wide association study (GWAS) involving 2.2 million individuals(*27*), we found genetic variants at the *FBLN2* locus are significantly associated with estimated glomerular filtration rate (eGFR), supported by methylation quantitative trait locus data(*28*), and single-nucleus ATAC-seq (snATAC-seq) (Figure 2K). In contrast, *FBLN1* showed no significant association with eGFR and *FBLN7* only weak association (Supplemental Figure 2, E and F). Together, these findings identify FBLN2 as a SMO-dependent ECM factor linked to CKD.

**Figure 2:**
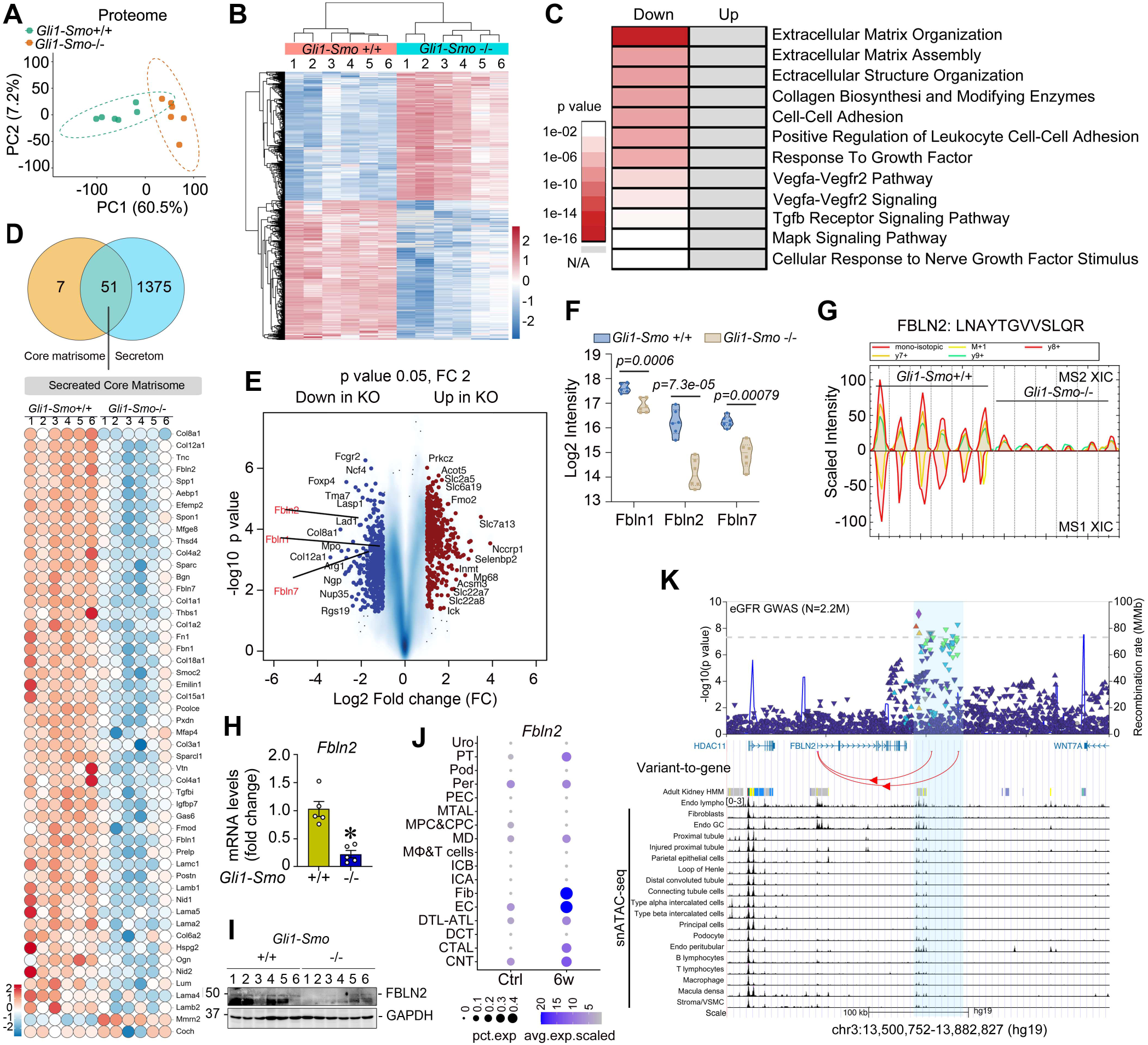
Global proteomics identifies FBLN2 as a prominent downstream matrisome component. In *Gli1-Smo*+/+ and *Gli1-Smo*-/- kidneys, after UIRI + UNx, (**A**) principal component analysis (PCA) of global proteomes; (**B**) Heatmap of significant proteins; label-free quantitation (LFQ) intensity was z-scored and plotted, highlighting two distinct protein clusters with varying abundance patterns. (**C**) Gene Ontology (GO) terms associated with protein clusters are plotted with names and statistical significance. (**D**) Heatmap of the differentially expressed core matrisome and secretome proteins. (**E**) Volcano plot of differential proteins. (**F**) pairwise comparisons of FBLN1, FBLN2, and FBLN7 (n = 6). In box plots, the lower and upper whiskers represent the minimum and maximum values, respectively; the box indicates the first and third quartiles, with the median shown as the center line. (**G**) Extracted ion chromatogram (XIC) graphs from Spectronaut software for FBLN2. MS1 XIC indicates peptide data from the first mass spectrometer, and MS2 XIC indicates peptide data from the second mass spectrometer (MS/MS). (**H-J**) qPCR analysis of *Fbln2* mRNA (**H**) (n=5); western blotting (**I**) of PBLN2 in *Gli1-Smo*+/+ and *Gli1-Smo*-/- kidneys. (**J**) Data mining (GSE139107) showing fibroblasts and endothelial cells as primary FBLN2 cellular sources. (**K**) Genetic ScoreCard of GWAS variants in the gene FBLN2 locus. The panels from top to bottom show the eGFR GWAS mapped in 2.2 million individuals, genes, variant-to-gene associations based on methylation QTL, chromatin state in adult kidney, and open chromatin in kidney cell types profiled by snATAC-seq. Scale bar, 25 µm. Arrows indicate positive cells. * *P* < 0.05. Graphs are presented as means ± SEM. Differences among groups were analyzed using two-sided unpaired t-tests.

### *Smo* deletion enhances tubular FAO and amino acid metabolism

Pathway enrichment of DIA proteomics revealed significant upregulation of metabolic pathways in *Gli1-Smo*-/- kidneys, particularly FAO and amino acid metabolism (Figure 3, A and B; Supplemental Figure 3, A and B). Key metabolic genes, including *Cd36*, *Cpt2*, *Acox3*, *Acadl*, *Glul*, *Gucy2d*, *Glud1*, *Slc25A21*, and *Slc25A44*, were elevated (Figure 3C), validated by western blotting (Figure 3D; Supplemental Figure 3C). CPT1α immunohistochemistry confirmed increased FAO in tubules (Figure 3E). ATP levels were higher (Figure 3F), whereas glycolysis remained largely unchanged (Supplemental Figure 3, D and E), indicating a selective shift toward oxidative metabolism. Moreover, several solute carriers were also upregulated, supporting substrate transport for enhanced metabolism (Figure 3G).

**Figure 3:**
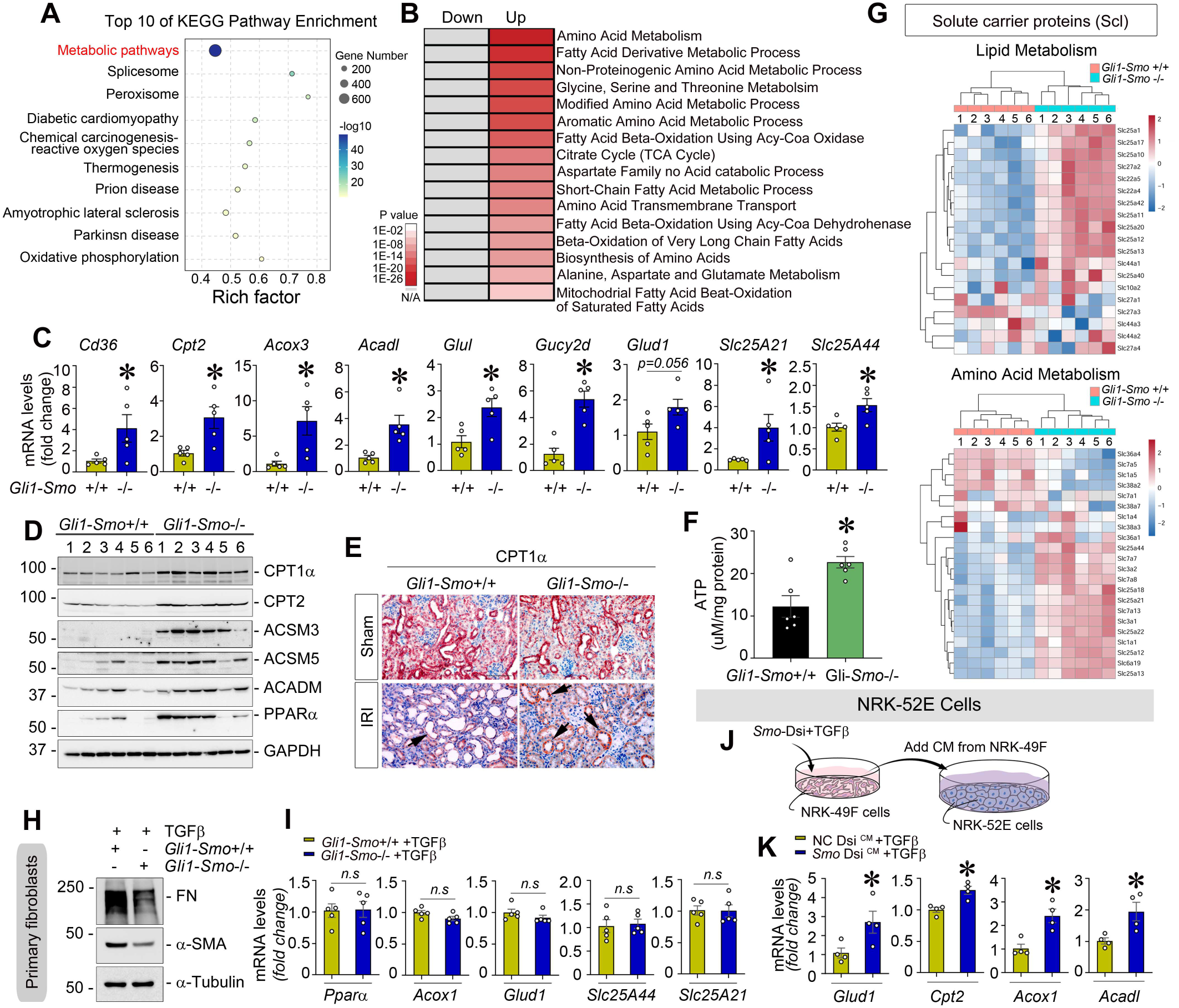
Loss of *Smo* in fibroblast enhances amino acid metabolism and fatty acid oxidation. In *Gli1-Smo*+/+ and *Gli1-Smo*-/- kidneys, after UIRI combined with nephrectomy, (**A**) Kyoto Encyclopedia of Genes and Genomes (KEGG) pathway enrichment analysis highlights upregulated metabolic pathways in fibrotic kidneys from both groups; (**B**) GO terms associated with protein clusters are plotted with names and statistical significance; (**C**) qPCR analysis showing *Cd36, Cpt2, Acox3*, *Acadl, Glul, Gucy2d, Glud1, Slc25A21,* and *Slc25A44* mRNA levels (n = 5); (**D**) western blotting demonstrating CPT1α, CPT2, ACSM3, ACSM5, ACADM, and PPARα protein level; (**E**) immunohistochemistry for CPT1α; (**F**) ELISA showing ATP levels in total kidneys (n = 6); (**G**) Heatmap of the differentially expressed solute carrier proteins in the amino acid metabolism and fatty acid oxidation (FAO) pathway for both groups after IRI. (**H, I**) Western blotting demonstrating the levels of FN and α-SMA (**H**), qPCR analysis of *Pparα*, *Acox1, Glud1, Slc25A44,* and *Slc25A21* (n = 5) (**I**) in primary cultured fibroblasts isolated from *Gli1-Smo*+/+ and *Gli1-Smo*-/- kidneys under TGFβ (2 ng/ml) for 24 hours. (**J**) Experimental design. (**K**) qPCR of *Glud1, Cpt2, Acox1,* and *Acadl* mRNA after incubated with *Smo*-deficient conditioned medium (CM) under TGFβ (2ng/ml) stimulation for 24 hours (n = 4). Arrows indicate positive staining. Scale bar, 25 µm. * *P* < 0.05. Graphs are presented as means ± SEM. Differences among groups were analyzed using two-sided unpaired t-tests.

Because *Smo* deletion was restricted to fibroblasts in this study, we next examined whether metabolic changes occurred within fibroblasts themselves. Primary kidney fibroblasts isolated from *Smo*-deficient mice exhibited reduced fibrotic activation under TGFβ stress, evidenced by decreased FN and α-SMA+ myofibroblasts activation (Figure 3H), but metabolic genes have no changes (Figure 3I), suggesting that the observed metabolic reprogramming occur in other kidney cell types. Given that tubular epithelial cells rely heavily on FAO for ATP generation, we hypothesized that fibroblast *Smo* deletion may influence tubular metabolism through intercellular communication. Indeed, normal rat kidney proximal tubular cells (NRK-52E) cultured with conditioned medium (CM) from *Smo*-knockdown fibroblasts (Figure 3J) displayed increased expression of FAO and amino acid metabolism genes (Figure 3K), indicating that fibroblast-derived factors regulate tubular metabolism.

### Acetyl-CoA acetyltransferase 1 (ACAT1) functions as a metabolic node linking FAO to amino acid utilization

Cross-analysis of the pathways of FAO and amino acid metabolism identified seven commonly upregulated mitochondrial enzymes, including IVD, ACAT1, AUH, ETFA, ETFB, ECHS1, and HSD17B10 (Figure 4, A and B). Among them, ACAT1 occupies a central position linking FAO, branched-chain amino acid catabolism, and ketone metabolism(*29*), was markedly increased in *Gli1-Smo-/-* kidneys (Figure 4, C-E; Supplemental Figure 4A). Immunostaining confirmed ACAT1 upregulation upon *Smo* loss and it was predominantly localized in tubular epithelial cells (Figure 4F), consistent with reanalysis in mouse snRNA-seq (GSE139107; Figure 4G). The snATAC analysis across 30 human kidney samples showed high open chromatin accessibility at the ACAT1 promoter in tubular cells(*27*), supporting active transcription in this compartment (Figure 4H). Consistently, ACAT1 expression in human proximal tubule segments decreased significantly (p < 1×10^-5^) in human AKI and CKD kidneys(*30*) (Supplemental Figure 4B), validated by immunostaining using human kidney biopsy specimens across multiple CKD etiologies, including IgA nephropathy, focal segmental glomerulosclerosis, and membranous nephropathy, compared with healthy controls (Figure 4I). Functionally, knockdown of *Acat1* with Dicer-substrate small interfering RNA (Supplemental Figure 4C) in tubular cells suppressed gene expression related to amino acid metabolism (*Glud1*, *Slc25A44*) and FAO (*Cpt2*, *Ppara*, *Acox1*, and *Acadl*) (Figure 4J) while enhancing fibrotic responses (Figure 4K), suggesting ACAT1 as a key metabolic checkpoint restraining fibrosis.

**Figure 4:**
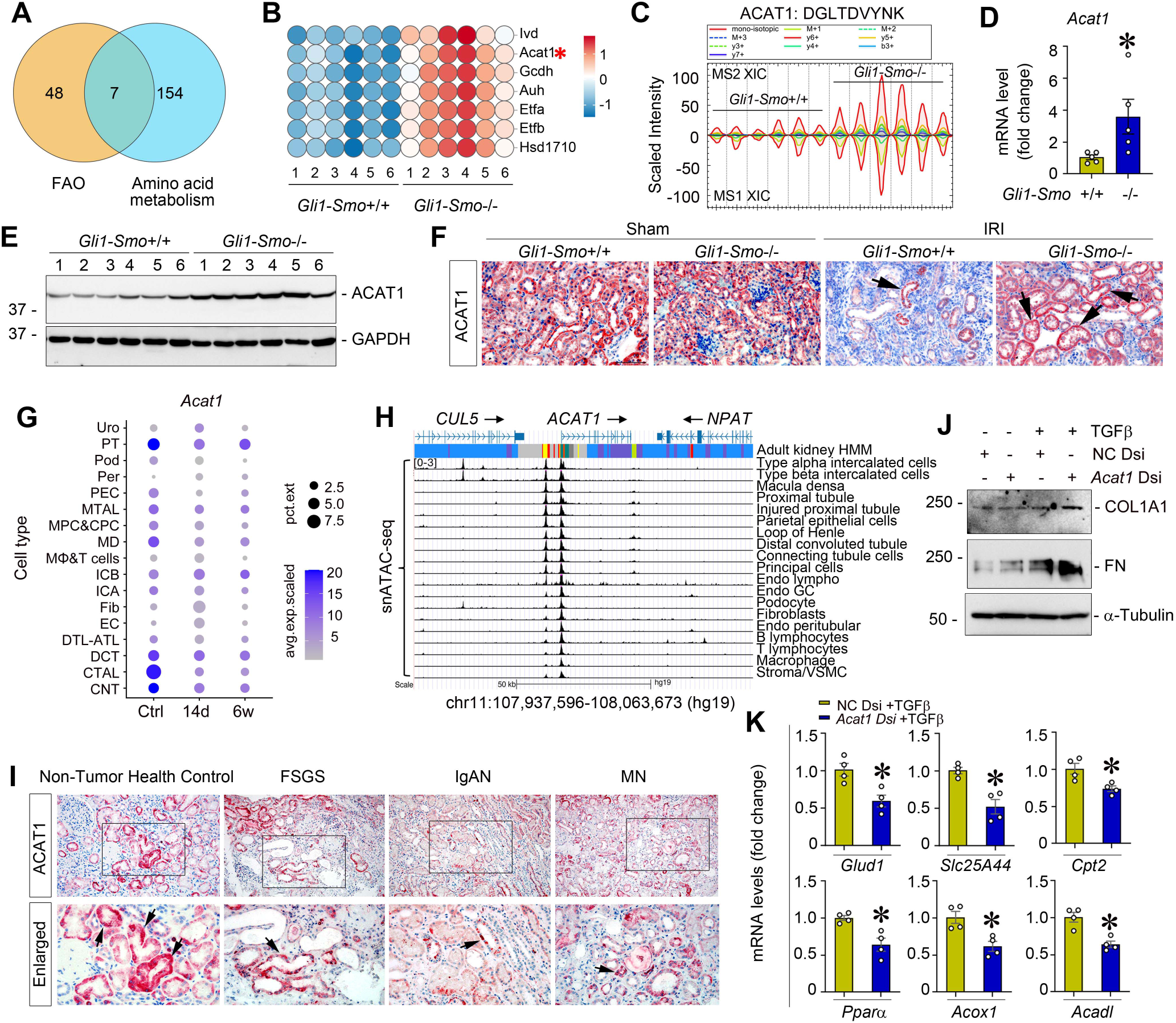
ACAT1 is a common regulatory node links amino acid metabolism to FAO. (**A**) Pie chart showing the distribution of significantly altered proteins and (**B**) heatmap of seven differentially expressed proteins across amino acid metabolism and FAO between *Gli1-Smo*+/+ and *Gli1-Smo*-/- kidneys after ischemic injury. (**C**) Extracted ion chromatogram (XIC) graphs from Spectronaut software for ACAT1. MS1 XIC indicates peptide data from the first mass spectrometer, and MS2 XIC indicates peptide data from the second mass spectrometer (MS/MS). (**D-F**) qPCR of *Acat1* mRNA(n=5) (**D**); western blotting (**E**) and immunostaining (**F**) of ACAT1 protein in *Gli1-Smo*+/+ and *Gli1-Smo*-/- kidneys. (**G**) Data mining (GSE139107) showing tubular cells as primary ACAT1 cellular sources. (**H**) snATAC-seq showing open chromatin in kidney tubular cells. (**I**) Immunohistochemical staining showing ACAT1 expression in kidney biopsy specimens from CKD patients. (**J, K**) qPCR of *Glud1, Slc25A44, Cpt2, Pparα, Acox1,* and *Acadl* mRNA (n = 4) (**J**), and western blot of COL1A1 and FN proteins **(K)** after transfected with Dicer-substrate *Acat1* siRNA under TGFβ (2ng/ml) stimulation for 24 hours. Arrows indicate positive staining. Scale bar, 25 µm. * *P* < 0.05. Graphs are presented as means ± SEM. Differences among groups were analyzed using two-sided unpaired t-tests.

### FBLN2 engages EGFR-AKT signaling as a functional ligand to regulate ACAT1

To investigate whether FBLN2 might be associated with metabolic alteration, we performed DIA-phosphoproteomics using the same set of kidney samples. PCA clearly separated *Gli1-Smo*+/+ and *Gli1-Smo*-/- kidneys (Figure 5A). In total, 26,923 phosphosites from 5,380 proteins were identified measured across samples. Correlations among biological replicates and intensity distributions are shown in Supplemental Figure 5, A and B. Kinase enrichment analysis revealed multiple signaling pathways altered in *Gli1-Smo*-/- kidneys (Figure 5B). Pairwise comparisons showed markedly reduced phosphorylation of AKT1 at Ser122 and Ser129 together with increased phosphorylation of ACAT1 at Ser230 and Ser238 in *Gli1-Smo*-/- kidneys (Figure 5, C-E). Western blotting confirmed reduced AKT phosphorylation in *Gli1-Smo*-/- kidneys (Figure 5F; Supplemental Figure 5C). Because AKT is commonly activated downstream of receptor tyrosine kinases(*31*), alternation in its phosphorylation often reflect changes in upstream receptor-mediated signaling. Among candidate receptors, epidermal growth factor receptor (EGFR) is a prime regulator of AKT and is well-known to respond to ECM cues(*32*). Consistent with this possibility, western blotting revealed reduced EGFR phosphorylation (Tyr1068) of in *Gli1-Smo*-/- kidneys (Figure 5G; Supplemental Figure 5D).

**Figure 5.**
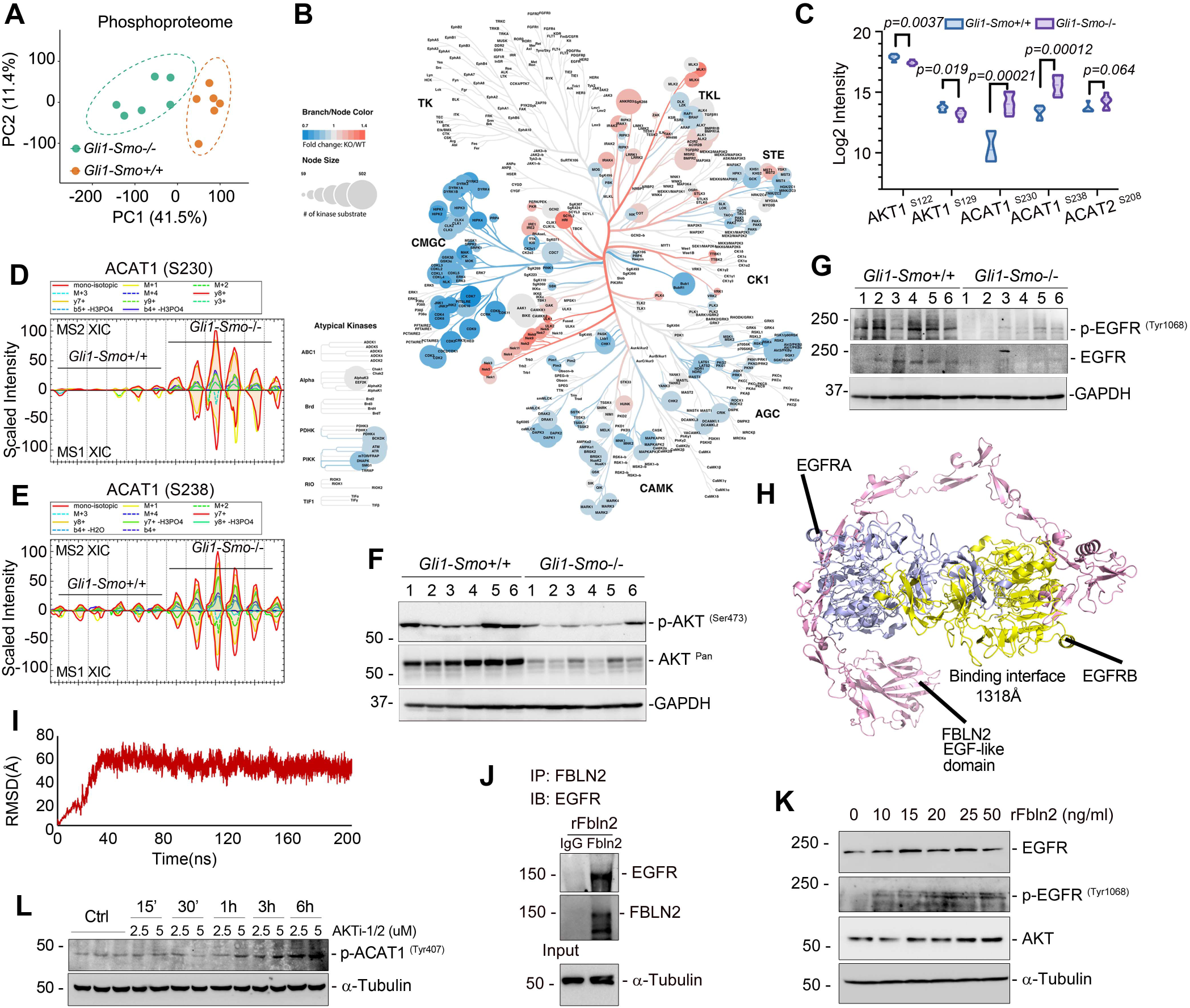
DIA-phosphoproteomics reveals EGFR–AKT inactivation following fibroblast *Smo* loss and identifies FBLN2 as a functional EGFR ligand. In *Gli1-Smo*+/+ and *Gli1-Smo*-/-kidneys, after UIRI + UNx, PCA of phosphoproteomes (**A**), poylogenetic tree of all protein kinases families detected (**B**), pairwise comparisons of AKT1 and ACAT1 (n = 5-6) (**C**), extracted ion chromatogram (XIC) graphs from Spectronaut software showing MS1 and MS2 phosphopeptide signals for ACAT1 (S230) **(D)** and ACAT1 (S238) **(E),** representing data from the first and second mass spectrometer (MS/MS), and western blot of p-AKT (Ser473), AKT **(F),** p-EGFR (Tyr1068), and EGFR (**G**). (**H**) Protein-protein docking analysis showing the two binding interfaces between FBLN2 and EGFR. (**I**) Molecular dynamics simulations showing FBLN2-EGFR complex rapidly stabilized after initial binding, reaching equilibrium within ∼30 ns and remaining stable throughout the simulation trajectory. (**J**) Immunoprecipitation revealing that FBLN2 physically binds to EGFR. NRK-52E cells treated with recombinant FBLN2 protein (rFBLN2) were immunoprecipitated with FBLN2 antibody, followed by immunoblotting with antibody against EGFR. (**K**) Western blotting showing p-EGFR (Tyr1068) and EGFR in NRK-52E cells after treatment with rFBLN2 at different dosages. (**L**) Western blotting of p-ACAT1 (Tyr407) in NRK-52E cells treated with AKTi-1/2. * *P* < 0.05. Graphs are presented as means ± SEM. Differences among groups were analyzed using two-sided unpaired t-tests.

Next, we asked whether FBLN2 could functionally engage EGFR. As a ∼130-kDa secreted ECM glycoprotein, FBLN2 contains 11 calcium-binding EGF-like domains that structurally resemble EGFR. These modules harbor conserved cysteine motifs that form disulfide-stabilized folds arranged a rigid bead-like structure coordinated by Ca²⁺, suggesting the potential to interact with the extracellular region of EGFR(*33*). To test this hypothesis, we performed structural modeling using AlphaFold3 and HADDOCK docking simulation, which predicted that FBLN2 can simultaneously contact two EGFR extracellular domains, stabilizing receptor dimerization (Figure 5H). Molecular dynamics simulations further indicated that the predicted FBLN2-EGFR complex rapidly stabilized after initial binding, reaching equilibrium within ∼30 ns and remaining stable throughout the simulation trajectory (Figure 5I). Hotspot analysis identified nine key residues such as E718, E607, D606, D604, Y746, and C941/Y960 at the FBLN2-EGFR interface (Supllemental Figure 5, E and F). These residues formed three interaction networks consisting of salt-bridge, guanidinium-mediated contacts, and aromatic-alkyl anchoring that collectively contributed substantially to the predicted binding free energy. These structural analyses support a model in which FBLN2 acts as a matrix-derived non-canonical ligand capable of promoting EGFR dimerization and activation. To experimentally validate this interaction, we performed co-immunoprecipitation assays. NRK-52E tubular cells treated with rFBLN2 protein resulted in robust co-precipitation of FBLN2 with EGFR, confirming their physical association (Figure 5J). Moreover, rFBLN2 induced EGFR phosphorylation at Tyr1068, demonstrating functional activation of EGFR-AKT signaling (Figure 5K). Because phosphoproteomics revealed increased ACAT1 phosphorylation at Ser230 and Ser238 in kidneys with reduced EGFR–AKT signaling, we next asked whether ACAT1 is responsive to perturbation of this signaling network. NRK-52E tubular cells were treated with the AKT inhibitor VIII (AKTi-1/2) across multiple doses and time points. AKT inhibition increased phosphorylation of ACAT1 at Tyr407, a regulatory site previously linked to ACAT1 tetramerization and activity in growth factor/tyrosine kinase signaling contexts. Since AKT is a serine/threonine kinase, this suggests that ACAT1 regulatory phosphorylation is indirectly coupled to the EGFR–AKT signaling state, potentially through the crosstalk with tyrosine kinase/phosphatase pathways (Figure 5L).

### Spatial lipidomics reveals altered fatty acid metabolism upon *Smo* Loss in fibroblasts

As a mitochondrial thiolase catalyzing the final thiolytic step of FAO, ACAT1 serves as a critical metabolic node linking fatty acid flux to broader lipid remodeling. To determine how fibroblast *Smo* deletion influences lipid metabolism in situ, we performed MALDI/MSI-based spatial lipidomics (Figure 6A). Unsupervised clustering revealed distinct lipid co-regulation patterns across samples (Figure 6B), while spatial segmentation separated cortex and medulla regions with clear genotype-dependent differences (Figure 6C). Across both compartments, 75 lipid species were detected and quantified (Methods and Supplementary Table S1), encompassing fatty acylcarnitines, glycerophospholipids, lysophospholipids, sphingolipids, and neutral lipids. Among these, 36 species were differentially regulated in the medulla and 29 in the cortex, with 22 shared alterations between both regions (Figure 6D).

**Figure 6.**
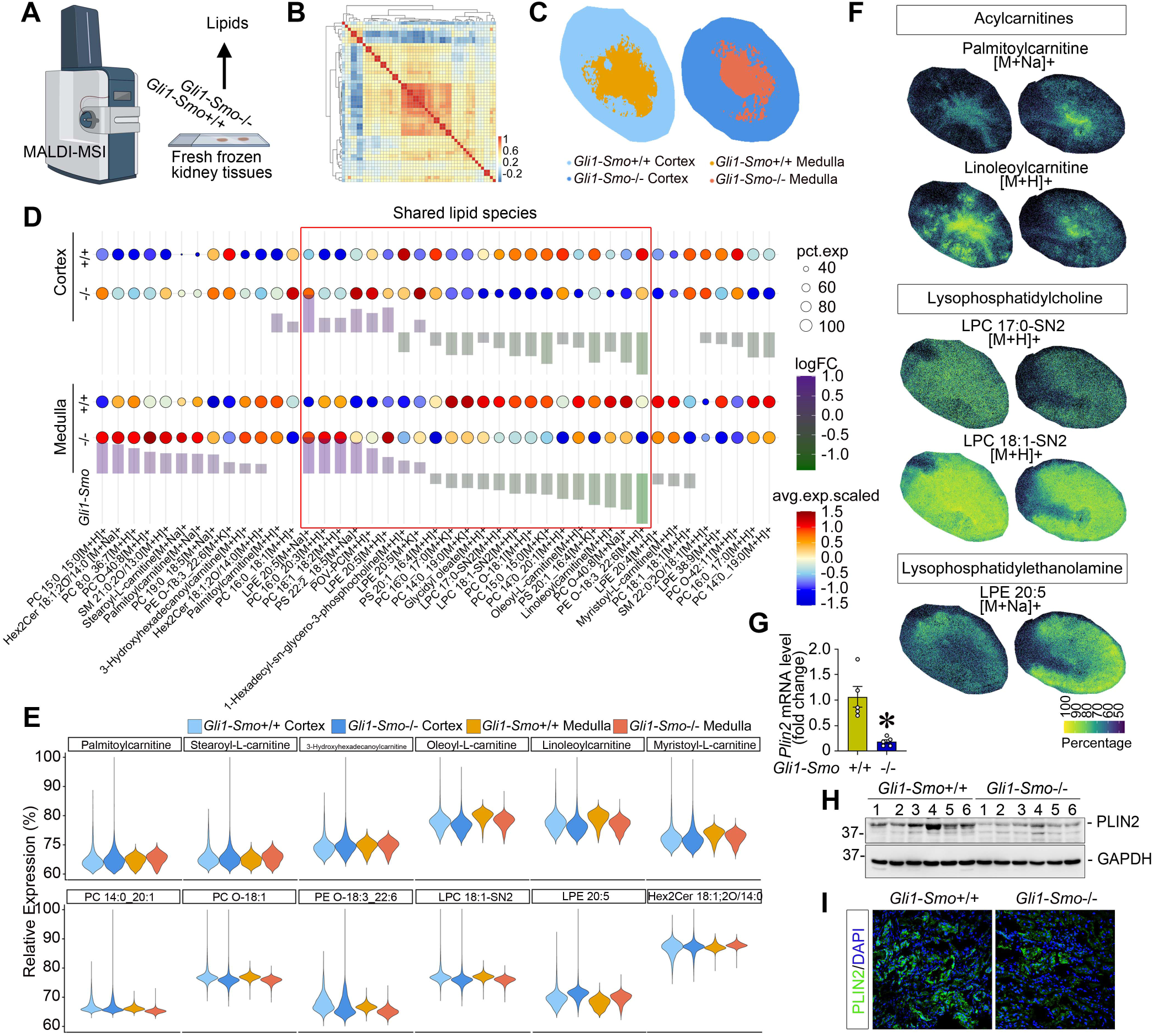
Spatial lipidomics reveals tubular metabolic remodeling after fibroblast-specific *Smo* deletion. (**A**) Schematic workflow of kidney tissue preparation and MALDI-MSI analysis of lipids from fresh-frozen *Gli1-Smo*+/+ and *Gli1-Smo*-/- kidneys. (**B**) Heatmap showing correlation of spatial lipid distributions across cortex and medulla regions in *Gli1-Smo*+/+ and *Gli1-Smo*-/-kidneys. (**C**) Representative segmentation maps of cortex and medulla based on lipid profiles. (**D**) Dot plot showing differentially expressed lipid species in cortex and medulla regions. (**E**) Violin plots showing relative abundance of lipid classes, including fatty acylcarnitines, phosphatidylcholines (PCs), lysophosphatidylcholines (LPCs), phosphatidylethanolamines (PEs), lysophosphatidylethanolamines (LPEs), and ceramides, in cortex and medulla of *Gli1-Smo*+/+ and *Gli1-Smo*-/- kidneys. (**F**) spatial ion images for representative lipid species in acylcarnitines, LPCs, and LPEs classes. (**G, H**) qPCR of *Plin2* mRNA (n=5) (**G**), and western blot of PLIN2 protein (**H**in *Gli1-Smo*+/+ and *Gli1-Smo*-/- kidneys after UIRI + UNx. (**I**) Immunofluorescence staining of PLIN2 (green) in *Gli1-Smo*+/+ and *Gli1-Smo*-/- kidneys after UIRI + UNx. Scale bar, 25 µm. DAPI is used as a nuclear counterstain. * *P* < 0.05. Graphs are presented as means ± SEM. Differences among groups were analyzed using two-sided unpaired t-tests.

Spatial lipidomic profiling revealed selective remodeling of FAO-linked acylcarnitine metabolism in *Gli1-Smo*-/- kidneys. Long-chain saturated acylcarnitines, including palmitoylcarnitine and stearoylcarnitine, were increased predominantly in the medulla, together with elevated 3-hydroxyhexadecanoylcarnitine, suggesting compartment-specific adaptation of mitochondrial FAO flux and intermediate handling. In contrast, unsaturated acylcarnitines, including oleoyl-L-carnitine and linoleoylcarnitine, were consistently reduced in both cortex and medulla, indicating decreased accumulation of lipotoxic incomplete oxidation products commonly associated with inflammation and fibrosis (Figure 6E, upper panel). Concomitantly, phosphatidylcholine (PC) species showed remodeling, including increased PC16:0_20:3 and decreased PC14:0_20:0 as well as ether-linked PCs (e.g., PC O-18:1), consistent with altered membrane lipid composition and remodeling dynamics. Lysophospholipids showed divergent regulation across classes: lysophosphatidylcholines (LPCs), including LPC 17:0-SN2 and LPC 18:1-SN2, were reduced, whereas lysophosphatidylethanolamine (LPE 20:5) was increased in *Gli1-Smo*-/- kidneys. Given that LPCs are bioactive lipids generated during phospholipid turnover and are implicated in pro-inflammatory and pro-fibrotic signaling(*34, 35*), their reduction suggested attenuation of inflammatory lipid signaling. In contrast, increased EPA-containing LPE 20:5 may reflect adaptive incorporation of omega-3-derived lipid substrates associated with anti-inflammatory and pro-resolving functions(*36*). In parallel, sphingolipid remodeling displayed strong spatial compartment specificity. Medullary levels of the oxidized glycosphingolipid Hex2Cer 18:1;2O/14:0 were increased (Figure 6E, lower panel), consistent with activation of glycosphingolipid pathways under oxidative and metabolic stress. Although glycosylated ceramides are generally considered less lipotoxic than free ceramides(*37*), their accumulation in oxidized form suggested stress-adaptive membrane remodeling rather than simple detoxification. Elevated oxidized phospholipids, such as POV-PC, further indicated compartmentalized accumulation of lipid oxidation products within discrete spatial domains (Figure 6D). Spatial ion maps for representative lipid species confirmed these region-specific alterations (Figure 6F).

Functionally, to measure lipid storage capacity, to assess lipid storage capacity, we evaluated perilipin 2 (PLIN2), a lipid droplet-associated protein and established marker of lipid accumulation(*13*). In *Gli1-Smo*-/- fibrotic kidneys, PLIN2 expression was markedly reduced at both mRNA and protein levels, and immunofluorescence staining confirmed decreased tubular Plin2 abundance (Figure 6, G-I; Supplemental Figure 6A). Notably, even during the early post-injury phase, when lipid metabolism has not yet become the dominant driver of tissue repair, we also observed increased expression of FAO-related proteins together with reduced lipid accumulation upon *Smo* loss, assessed by Oil Red O staining (Supplemental Figure 6, B-E).

### Pdgfrβ+ fibroblast *Smo* deletion independently validates the metabolic-fibrotic axis

Considering the heterogeneity of kidney fibroblast populations, we next sought to validate our findings in an independent Pdgfrβ+ fibroblast lineage. To this end, we generated tamoxifen-inducible fibroblast-specific *Smo* knockout mice by crossing *Pdgfrβ-P2A-CreER^T2^* with *Smo*-floxed mice. Following tamoxifen induction, Pdgfrβ-Smo-/- mice were subjected to ischemic injury and UUO (Figure 7A). Consistent with the phenotype observed in *Gli1-Smo*-/- mice, *Pdgfrβ-Smo*-/- mice exhibited preserved Scr and BUN levels after ischemic injury (Figure 7B), accompanied by reduced fibrosis and inflammation in both models (Figure 7, C-F; Supplemental Figure 7, A-D), as assessed by qPCR, western blotting, and histological analyses. These protective effects were associated with decreased FBLN2 expression, alongside enhanced amino acid metabolism and FAO, including upregulation of ACAT1 and related metabolic enzymes (Figure 7, G-I; Supplemental Figure 7E). In parallel, EGFR-AKT signaling was suppressed in *Pgdfrβ-Smo*-/- kidneys (Figure 7J; Supplemental Figure 7F). Consistent with this metabolic shift, lipid storage was markedly reduced, as evidenced by decreased PLIN2 expression and spatial distribution (Figure 7, K-M; Spplemental Figure 7G). These findings therefore independently validated and generalized the fibroblast SMO-tubular metabolic axis across distinct fibroblast lineages, linking fibroblast SMO signaling to ECM-dependent regulation of epithelial lipid metabolism.

**Figure 7.**
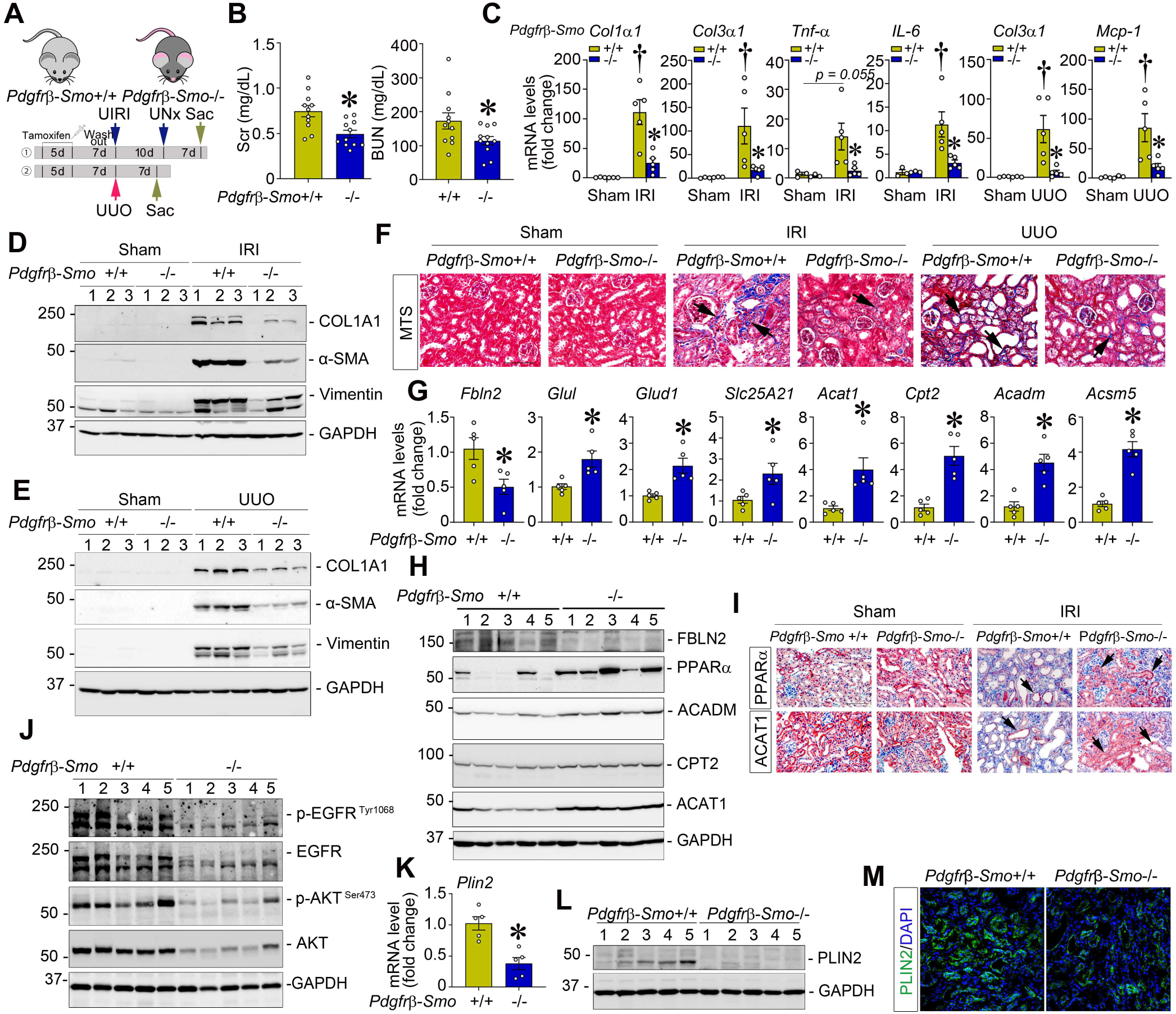
Loss of *Smo* in *Pdgfrβ* + fibroblasts mitigate kidney fibrosis through the FBLN2-EGFR-ACAT1 metabolic-fibrotic axis. (**A**) Experiment design. *Pdgfrβ-Smo*+/+ and *Pdgfrβ-Smo*-/- mice were administered tamoxifen prior to UIRI combined with nephrectomy (model ①) or UUO (model ②). In model ①, UNx after UIRI, and mice were sacrificed 7 days thereafter. In model ②, mice were sacrificed 7 days after UUO. In *Pdgfrβ-Smo*+/+ and *Pdgfrβ-Smo*-/- kidneys, after UIRI + UNx or UUO, (**B**) Scr and BUN levels (n = 11), (**C**) qPCR of *Col1a1*, *Col3a1*, *Tnf-a, IL-6,* and *Mcp-1* mRNA (n = 5; Sham, n = 3), (**D, E**) Western blotting of COL1A1, α-SMA, and Vimentin proteins, (**F**) Representative micrographs of MTS and immunohistochemical staining for Vimentin, COL1A1, and CD45, (**G**) qPCR analysis showing *Fbln2*, *Glul, Glud1, Slc25A21, Acat1, Cpt2, Acadm,* and *Acsm5* levels (n = 5), (**H**) western blotting of FBLN2, PPARα, ACADM, CPT2, ACAT1, (**I**) immunohistochemistry for PPARα and ACAT1, (**J**) western blotting of p-EGFR (Tyr 1068), EGFR, p-AKT (Ser473), and AKT, (**K**) qPCR of *Plin2* mRNA (n=5), (**L**)western blotting of PLIN2 protein, and (**M**) immunofluorescence staining of PLIN2 (green). Scale bar, 25 µm. Arrows indicate positive cells. DAPI is used as a nuclear counterstain. † *P* < 0.05 versus sham control, * *P* < 0.05 versus *Pdgfrβ-Smo*+/+ mice after UIRI+UNx or UUO. Graphs are presented as means ± SEM. Differences among groups were analyzed using two-sided unpaired t-tests or one-way ANOVA followed by the Student-Newman-Keuls test.

### FBLN2 mediates fibroblast-tubular crosstalk to regulate EGFR-AKT signaling and ACAT1-dependent metabolism in vitro and ex vivo

Having established this axis *in vivo*, we next sought to delineate the underlying fibroblast-epithelial signaling mechanism. In normal rat kidney fibroblasts (NRK-49F), genetic knockdown of *Smo* using Dicer-substrate siRNA or pharmacological inhibition of *Smo* with cyclopamine (CPN) reduced expression of fibrosis-related proteins under TGF-β stress and suppressed FBLN2 expression and (Figure 8, A-C), without affecting AKT signaling within fibroblasts themselves (Figure 8, B and D). Conversely, rFBLN2 enhanced fibroblast activation but similarly did not alter AKT expression (Figure 8D), suggesting that FBLN2 primarily acts on neighboring epithelial cells rather than fibroblasts.

**Figure 8.**
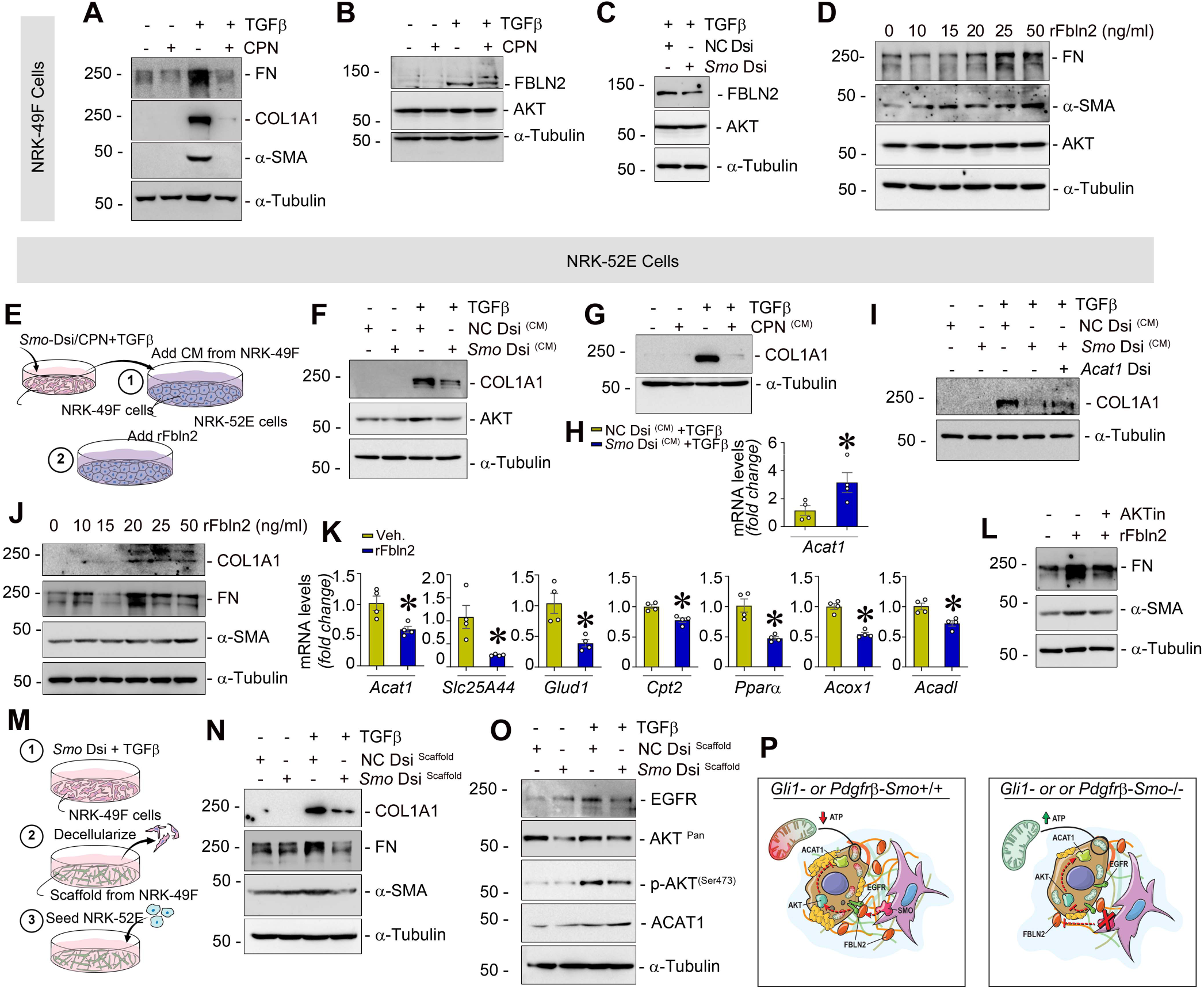
Fibroblast Smo controls FBLN2-dependent fibroblast–tubular crosstalk to modulate EGFR–AKT signaling and ACAT1 metabolism. (**A-D**) NRK-49F cells treated with Smo inhibitor cyclopamine (CPN, 2.5 µM) (**A, B**), transfected with NC- or *Smo*-Dsi (**C**) under TGFβ stress or rFBLN2 at different dose (**D**) for 24 hours. Western blotting showing the protein expression of FN, COL1A1, α-SMA, FBLN2, and AKT. (**E**) Experimental design. (**F-H**) Western blotting of COL1A1 and AKT proteins (**F, G**), and qPCR of *Acat1*(**H**) in NRK-52E cells after incubated with FBLN2-deficient conditioned medium (CM). (**I**) Western blotting showing that FBLN2-deficient CM reduced the induction of COL1A1 under TGFβ stress, but they were increased after knockdown of *Acat1*. (**J**) Western blotting of COL1A1, FN, and α-SMA after rFBLN2 treatment for 24 hours. (**K**) qPCR of *Acat1, Slc25A44, Glud1, Cpt2, Pparα, Acox1*, and *Acadl* after treated with rFbln2 (n=4). (**L**) Western blotting of FN and α-SMA after rFBLN2 and AKT inhibitor treatment for 24 hours. (**M**) Schematic diagram. (**N-O**) Western blots of COL1A1, FN, α-SMA (**N**), EGFR, AKT, p-AKT (Ser473), and ACAT1(**o**) in NRK-52E cells seeded on FBLN2-deficient matrix scaffold isolated from *Smo*-knockdown fibroblasts under TGFβ stress. (**p**) Diagram illustrating loss of fibroblast-derived *Smo* inhibited FBLN2 to interacts with EGFR/AKT signaling pathway to active tubular lipid metabolic reprogramming through ACAT1 to determine CKD outcomes. * *P* < 0.05. Graphs are presented as means ± SEM. Differences among groups were analyzed using two-sided unpaired t-tests.

To model fibroblast-tubular communications, NRK-52E tubular epithelial cells were exposed to CM from *Smo*-knockdown/CPN-treated fibroblasts or incubated with increasing doses of rFBLN2 (Figure 8E). CM from *Smo*-deficient fibroblasts suppressed COL1A1 and AKT protein (Figure 8, F and G) while increasing *Acat1* transcription (Figure 8H) in tubular cells. Notably, *Acat1* knockdown abolished the reduction in COL1A1 induced by CM from *Smo*-deficient fibroblasts (Figure 8I), demonstrating that ACAT1 is required for the antifibrotic metabolic response. In contrast, rFBLN2 treatment increased COL1A1, FN, and α-SMA (Figure 8J). It suppressed amino acid metabolism and FAO related genes, including *Acat1, Slc25A44, Glud1, Cpt2, Pparα, Acox1*, and *Acadl* (Figure 8K). Pharmacological inhibition of AKT with AKTi-1/2 blocked rFBLN2-induced FN and α-SMA expression (Figure 8L).

To better recapitulate *in vivo* microenvironment, we further performed *ex vivo* experiments using decellularized scaffolds isolated from fibroblasts under TGF-β stimulation, which were reseeded with NRK-52E cells, a system well-established in our lab(*21*) (Figure 8M). Tubular epithelial cells cultured on these *Smo*-deficient fibroblast-derived scaffolds exhibited reduced COL1A1, FN, α-SMA, EGFR, AKT, and p-AKT (Ser473) levels, accompanied by increased ACAT1 expression (Figure 8, N and O). Collectively, these findings establish fibroblast-epithelial pipelines in which SMO-dependent secretion of FBLN2 activates EGFR-AKT signaling in tubular epithelial cells, suppressing ACAT1-driven oxidative metabolism and promoting fibrotic remodeling (Figure 8P).

## Discussion

Across organs, fibrosis represents the final common pathway of chronic injury and the tipping point from adaptive repair to irreversible organ dysfunction. Rather than reflecting simple ECM accumulation, our findings suggested that fibrotic progression is driven by disrupted metabolic coordination between fibroblasts and epithelial compartments after CKD. In this model, the ECM is more than structural scaffolding, it is a dynamic signaling interface that shapes cellular bioenergetics. Fibrosis therefore emerges when matrix-dependent metabolic coupling fails, locking tissue into maladaptive remodeling.

A key conceptual advance of this study is that fibroblast-intrinsic SMO functioned as a signal integrator that coupled ECM spectrum remodeling to tubular lipid metabolic reprogramming, thereby dictating fibrotic trajectory. As a GPCR-like Hedgehog receptor, SMO is classically associated with developmental patterning and Gli-dependent transcription(*38*). However, GPCR-like receptors possess broader capacities: they integrate extracellular stress signals, tune kinase cascades, and orchestrate metabolic adaptation(*39, 40*). Our work extended SMO’s functional scope beyond development, positioning it as a regulator of matrix architecture and cross-compartmental metabolic coordination in adult kidney injury.

Although Hedgehog signaling has been implicated in fibroblast activation, prior models have largely focused on expansion of myofibroblasts(*41–44*). Our study showed that SMO governs the ECM itself (Fig. 2). During acute injury, loss of Smo in fibroblasts transiently activated matrix proteins such as Nidogen 1, facilitating protective crosstalk with tubular Wnt signaling(*22*). In contrast, chronic perturbation profoundly reshaped matrix reassembly and spectrum (Fig. 2), suggesting that SMO controlled not only matrix quantity but also its signaling competence. Mechanistically, we speculated that SMO tunes the biochemical and spatial properties of the interstitial niche. By tuning fibroblast secretory output, SMO defines which matrix components are exposed, organized, and available to engage epithelial receptors. In this way, SMO-dependent matrix remodeling becomes functionally instructive, establishing a microenvironment that propagates metabolic signals across compartments.

Among the core matrisome components influenced by SMO, FBLN2 emerges as a prominent mediator. Traditionally considered as a structural ECM glycoprotein contributing to elastic fiber assembly and matrix stabilization, our findings expanded its role: FBLN2 functioned as a previously unrecognized functional ligand or binding partner of EGFR in fibrotic kidneys (Fig. 5). Rather than relying solely on soluble growth factors, matrix-embedded FBLN2 provides a spatially organized platform for EGFR engagement. It physically interacts with EGFR (Fig. 5), facilitating its phosphorylation and mitochondrial translocation. This observation expands the conceptual role of FBLN2 from structural scaffolds to modulators of receptor trafficking and intracellular localization. Through the FBLN2-EGFR-AKT axis, ECM composition directly governs intracellular metabolic programs. Disruption of SMO-dependent FBLN2 organization selectively impaired EGFR phosphorylation and downstream AKT activation, thereby reshaping tubular metabolic flux, whereas AKT activity within fibroblasts themselves remained largely unaffected (Fig. 3). Notably, Smo knockdown in fibroblasts can affect its autophagy(*45*), suggesting that SMO may regulate distinct intracellular programs in fibroblasts while exerting paracrine metabolic effects on neighboring epithelial cells. This compartmental specificity reflects that fibroblast-derived matrix cues drive their pathogenic influence primarily through reprogramming epithelial bioenergetics rather than by reinforcing fibroblast-intrinsic activation programs.

The downstream metabolic consequences of this axis are substantial. Properly regulated EGFR-AKT signaling supported FAO and amino acid metabolism, pathways essential for maintaining mitochondrial function under fibrosis. When this signaling platform is disrupted, bioenergetic flexibility is compromised, lipid accumulation ensues, and maladaptive remodeling progresses. In this context, FBLN2 thus acts not merely as a structural ECM protein but as a ligand-like organizer that translates matrix composition into mitochondrial instruction. A newly identified effector of this cascade is ACAT1, a mitochondrial enzyme that catalyzing the reversible condensation of acetyl-CoA and regulating flux through FAO and ketogenesis pathways. While previously studied in cancer and systemic lipid metabolism(*46, 47*), ACAT1’s role in fibrosis was unexplored. Our findings positioned ACAT1 as a metabolic gatekeeper downstream of SMO-FBLN2-EGFR axis (Fig. 4). ACAT1 induction coincided with enhanced mitochondrial fatty acid oxidation signatures, indicating activation of oxidative metabolic programs in tubular epithelial cells. ACAT1 reprogramming altered lipid classes, including acylcarnitines, PCs, LPCs, and LPEs, consistent with enhanced mitochondrial lipid flux (Figure 6). These lipidomic shifts reinforce the notion that matrix remodeling is metabolically instructive. It likely impacts membrane dynamics, mitochondrial structure, and signaling microdomains, amplifying the metabolic consequences of upstream matrix cues. Surprisingly, current GWAS analyses do not directly associate ACAT1 with CKD susceptibility although its expression was markedly reduced in CKD biopsy specimens with different etiologies (Figure 4), highlighting a potential disconnect between inherited genetic risk and metabolically critical regulatory nodes. This absence represents an underappreciated metabolic function of ACAT1 that may not be captured by genetic association studies and deserving deeper exploration.

The clinical relevance of this study is further supported by human genetic data linking FBLN2 loci with GFR in CKD patients. The convergence of unbiased proteomics, in vivo genetic models, and human association analyses strengthens the translational significance of this matrix-metabolism axis. Moreover, SMO itself is pharmacologically tractable(*41, 48*), raising the prospect that selective modulation of fibroblast SMO could recalibrate matrix organization and restore metabolic coupling without broadly suppressing Hedgehog signaling.

Beyond kidney disease, these findings have broader implications. Fibroblasts in the heart, lung, and liver share conserved developmental programs and ECM secretory machinery. It is plausible that SMO-dependent matrix reorganization similarly governs metabolic adaptation in other fibrotic contexts. The SMO-FBLN2-EGFR-ACAT1 axis may therefore represent a generalizable paradigm in which developmental GPCR-like receptors coordinate structural and metabolic circuits to determine tissue fate.

Our findings also invite rethinking of intercellular communication during fibrosis. Conventional models emphasize soluble factors or direct receptor-ligand interactions. In contrast, our data supported a model in which matrix architecture itself serves as an intermediary signaling layer. By reorganizing ECM components, fibroblasts reshape the biochemical landscape that dictates receptor activation and mitochondrial function in neighboring cells. Fibrosis arises when this matrix-mediated metabolic dialogue becomes dysregulated.

In summary, our study identifies FBLN2 as a fibroblast-derived ECM effector that translates SMO-dependent matrix remodeling into tubular metabolic dysfunction during kidney fibrosis. By coupling FBLN2–EGFR–AKT signaling to suppression of ACAT1-dependent mitochondrial fatty acid and amino acid metabolism, this pathway provides a mechanism by which stromal matrix changes directly shape epithelial bioenergetics. These findings reposition kidney fibrosis as not only a disorder of matrix accumulation, but also a disease of disrupted stromal–epithelial metabolic communication. Targeting this matrix-metabolic crosstalk may offer a strategy to restore tubular metabolism and slow chronic kidney disease progression.

## Methods

### Study approval

All animal experiments were approved by the Institutional Animal Care and Use Committee of the University of Connecticut School of Medicine and were conducted in accordance with institutional and federal regulations for the humane care and use of laboratory animals. Studies involving human samples were approved by the Institutional Review Board of the University of Connecticut School of Medicine. Our CKD study exclusively examined male mice because female mice represented greater resistance than male mice against renal IRI and UUO. Human kidney biopsy samples were obtained from both men and women. It is unknown whether the findings in male mice are relevant to female mice.

### Mice

Homozygous *Smo*-floxed mice were generously gifted by Dr. Anna Mae Diehl at Duke University. *Gli1*-CreER^T2^ (#007913), and *Pdgfrβ*-P2A-CreER^T2^ (#030201) were purchased from The Jackson Laboratory. By mating *Smo*–floxed mice with *Gli1*-CreERT2 or *Pdgfr*β-P2A-CreERT2 transgenic mice, conditional knockout mice in which *Smo* gene was specifically disrupted in kidney Gli1+ or Pdgfrβ+ fibroblasts (genotype *Smo*^fl/fl^, Cre ±) were created. These mice were crossbred with homozygous *Smo*–floxed mice (genotype *Smo*^fl/fl^) to generate offspring with 50% *Gli1-Smo*-/- or *Pdgfr*β-Smo-/- mice and 50% control mice (*Gli1-Smo*+/+ or *Pdgfr*β-Smo+/+) within the same litters. A routine PCR protocol was used for genotyping of tail DNA samples with the following primer pairs: Cre transgene (*Gli1*-CreERT2), 5’-GCG-GTC-TGG-CAG-TAA-AAA-CTA-TC-3’ and 5’-GTG-AAA-CAG-CAT-TGC-TGT-CAC-TT-3’; Cre transgene (*Pdgfr*β-P2A-CreERT2), 5’-AGC-TTG-TGG-CAG-TGT-AGC-TG-3’, 5’-ACA-TGT-CCA-TCA-GGT-TCT-TGC-3’, and 5’-CCA-CCT-TGA-ATG-AAG-TCA-ACA-C-3’; and *Smo* genotyping, 5’-CGC-ACC-GGT-CGC-CTA-AGT-AGC-3’ and 5’-GGC-GCT-GTG-AGG-CCC-AGG-C-3’. All animals were fed regular chow (LabDiet, 5V5M-PicoLab Verified 50 IF/9F) and housed under standard condition (69° to 72° F, 12-hour light/dark cycle, and relative humidity of 40–60%) in the university animal facility.

### CKD models

Two murine CKD models were employed: unilateral IRI with contralateral nephrectomy and UUO. For unilateral IRI, with contralateral nephrectomy, the left renal pedicle was clamped for 35 minutes, with body temperature maintained between 36-37.5°C, the contralateral kidney was removed 10 days later, and mice were sacrificed 7 days thereafter. UUO was induced by unilateral ureteral ligation for 7 days.

### Biochemical and ELISA measurements

Serum BUN, and creatinine were measured using the QuantiChromTM Urea (DIUR-100), and Creatinine (DICT-500) assay kits (BioAssay Systems), respectively. Kidney ATP content was assessed using the ATP Colorimetric/Fluorometric Assay Kit (ab83355, Abcam).

### Human Kidney Biopsy Specimens

The blank sections of human kidney biopsy specimens were obtained from the University of Pittsburgh Medical Center. Nontumor kidney tissue from patients with renal cell carcinoma who underwent nephrectomy was used as normal controls. All patients in the presented study signed informed consent forms before they underwent kidney biopsy or nephrectomy.

### Histology and immunostaining

Paraffin-embedded kidney sections (3 μm) were stained with PAS and MTS following standard protocols. For immunohistochemical staining, sections were incubated with primary antibodies at 4°C overnight, followed by HRP-conjugated secondary antibody. Non-immune normal IgG served as a negative control. Slides were imaged using an Olympus BX43 microscope. For immunofluorescence, kidney cryosections were fixed with 3.7% paraformaldehyde for 15 min at room temperature, blocked with 10% donkey serum for 1 hour, and then incubated with primary antibodies at 4°C overnight. After washing, Alexa Fluor 488-conjugated secondary antibody was applied. Slides were imaged using an Olympus FluoView 1000 confocal microscope. Detailed antibody information is provided in Supplementary Table S2.

### qPCR

Total RNA was extracted using TRIzol reagent (Invitrogen) and reverse-transcribed into cDNA (BioRad). qPCR was performed on a CFX Connect Real-Time PCR detection system (Bio-Rad). Gene expression levels were normalized to β-actin. Primers sequences are listed in Supplementary Table S3.

### Western blot

Kidney tissues were lysed on ice with radioimmune precipitation assay buffer containing 1% NP-40, 0.1% SDS, 100 μg/ml PMSF, 1% protease inhibitor cocktail, and 1% phosphatase I and II inhibitor cocktail in PBS. Lysates were centrifuged at 13,000×g at 4°C for 15 min, and the supernatants were collected for Western blot. Detailed antibody information is provided in Supplementary Table S2.

### DIA-based global- and phospho-proteomics

Kidney tissues were processed using a previously published proteomics workflow(*49, 50*). In brief, mouse kidneys were lysed in SDS buffer (4% SDS, 50 mM EDTA, 20 mM DTT, 2% Tween-20, 100 mM Tris-HCl, pH 8.0) and homogenized by sonication using a Misonix Sonicator 3000 Ultrasonic Cell Disruptor at 4°C for 10 minutes (5-second on/off cycles) to ensure complete lysis. The lysates were then centrifuged at 20,000 × g for 1 hour to remove insoluble material. Approximately 800 μg of protein was used for subsequent digestion. Reduction and alkylation were performed with 10 mM dithiothreitol (DTT) for 1 hour at 56°C, followed by 20 mM iodoacetamide (IAA) in the dark for 45 minutes at room temperature. The reduced and alkylated proteins were then subjected to a precipitation-based digestion as described previously (*51*). Briefly, five volumes of precooled precipitation solution containing 50% acetone, 50% ethanol, and 0.1% acetic acid were added to the protein mixture and kept at −20 °C overnight. The mixture was centrifuged at 20,000 x g for 40 min. The precipitated proteins were washed with precooled 100% acetone and 70% ethanol, and centrifuged at 20,000 x g, 4 °C for 40 min. After final wash, the remaining solvent was evaporated in a SpeedVac for 5 min. Samples were then added with 300 μL 100 mM NH4HCO3 and digested overnight at 37°C with trypsin (Promega) at a 1:20 (w/w) enzyme-to-protein ratio. Digested peptides were purified using C18 columns (MacroSpin Columns, NEST Group Inc.). Five percent of the peptides was used for total proteome analysis, while the remaining 95% was used for phosphopeptide enrichment.

Phosphopeptide enrichment was performed using the High-Select Fe-NTA kit (Thermo Fisher Scientific, A32992) according to the manufacturer’s instructions(*52, 53*). Briefly, the resin from the spin columns was aliquoted and incubated with 250 μg of total peptides for 30 minutes at room temperature, then transferred to filter tips (TF-20-L-R-S, Axygen). The supernatant was removed by centrifugation. The resin-bound phosphopeptides were subsequently washed three times with 200 μL of wash buffer (80% acetonitrile, 0.1% trifluoroacetic acid), followed by two washes with 200 μL of water to remove nonspecifically bound peptides. Phosphopeptides were then eluted twice with 100 μL of elution buffer (50% acetonitrile, 5% NH₃·H₂O). All centrifugation steps were performed at 500 × g for 30 seconds. The eluates were dried in a SpeedVac and stored at −80°C prior to mass spectrometry (MS) analysis.

Samples were analyzed by data-independent acquisition (DIA) mass spectrometry as previously described(*54–56*). Measurements were performed on an Orbitrap Fusion Tribrid mass spectrometer (Thermo Scientific) coupled to a NanoFlex nanoelectrospray source and an EASY-nLC 1200 system. Peptides were separated using a 120-min gradient at 300 nL/min, with the column maintained at 60°C (PRSO-V1, Sonation GmbH). The DIA method included one MS1 scan followed by 33 MS2 scans with variable isolation windows and 1 m/z overlap. MS1 spectra were acquired from 350–1650 m/z at 120,000 resolution (m/z 200), with an AGC target of 2 × 10⁶ and 100 ms maximum injection time. MS2 spectra were acquired from 200–1800 m/z at 30,000 resolution, using 28% normalized HCD collision energy, an AGC target of 1.5 × 10⁶, and 50 ms maximum injection time. The default charge state was set to 2. Both MS1 and MS2 data were collected in profile mode.

Raw files were processed in Spectronaut v18(*57–59*), using the directDIA workflow against the SwissProt mouse FASTA database. Methionine oxidation was set as a variable modification and carbamidomethylation of cysteine as a fixed modification. For phosphoproteomics, phosphorylation (S/T/Y) was included as an additional variable modification. Peptide and protein FDRs were controlled below 1%. For phosphosite analysis, identifications were filtered with PTM score > 0.75, while quantitative tables were filtered with PTM score > 0.01 to reduce missing values caused by localization ambiguity.

### MALDI MSI-based Spatial Lipidomic

The kidney tissue was embedded in gelatin immediately after dissection from the mice. The samples were cryosectioned at 10 µm thickness, mounted onto ITO-coated slides, and stored at - 80 °C until analysis. Wild-type and knockout tissues were placed on the same slide to minimize experimental variation. Prior to matrix application, the tissue sections were dried at room temperature for 30 min. A 10 mg/mL HCCA (α-cyano-4-hydroxycinnamic acid) matrix was applied using an HTX TM-Sprayer M3+. The matrix solution was prepared in 70% acetonitrile containing 0.2% TFA. The spraying parameters were set as follows: nozzle temperature 75 °C, flow rate 120 µg/min, and 4 passes. No drying time was used between passes, and all other parameters were kept at the default settings.

The MALDI imaging experiments were performed on a Bruker timsTOF fleX MALDI-2 mass spectrometer. Instrument parameters were configured using timsControl, and MSI data acquisition was carried out with flexImaging software on positive mode. Data were acquired over an m/z range of 300 −1300 with an ion mobility (1/K₀) range of 0.66 - 1.74 V·s/cm² and a TIMS ramp time of 200 ms. The collision energy and RF amplitude were set to 10 eV and 1800 Vpp, respectively. The quadrupole ion energy was 4 eV. The pre TOF transfer time and pre pulse storage time were 85 µs and 10 µs, respectively. Imaging was performed at a spatial resolution of 20 µm with beam scan enabled, using 200 laser shots per pixel at a laser frequency of 10 kHz. Both m/z and ion mobility calibrations were performed prior to data acquisition using the Agilent ESI Low Concentration Tune Mix. The gas flow was adjusted to position the m/z 622 calibrant ion at 164 UV before ion mobility calibration. All other instrument parameters were kept at default settings.

### Lipid extraction for LC-MS/MS

The remaining kidney tissue used for MALDI-MSI was subjected to LC-MS/MS lipidomics to generate a reliable target lipid library for subsequent MSI data annotation. Tissue samples were snap-frozen in liquid nitrogen and mechanically disrupted using a Covaris CP02 CryoPREP Automated Dry Pulverizer. The pulverized tissue was transferred to 2 mL tubes containing water and lysed by sonication at 4 °C for 10 cycles (1 min per cycle) using a VialTweeter device (Hielscher Ultrasound Technology). For lipid extraction, three volumes of pre-chilled methanol were added to the tissue lysate and vortexed for 10 s. Subsequently, seven volumes of pre-chilled methyl tert-butyl ether (MTBE) were added, and the samples were shaken for 1 h at room temperature. Finally, two volumes of pre-chilled water were added into the sample, followed by vortexing for 10 s. The samples were then centrifuged at 1,000 × g for 5 min to separate the phases. The upper organic phase was collected and transferred to a new tube and dried using a SpeedVac concentrator. The dried lipid extracts were reconstituted in 200 µL of BuOH:IPA:H₂O (8:23:69, v/v/v)(*60*), and 1 µL of the lipid was injected for LC-MS/MS analysis.

### LC-MS/MS lipidomics for library generation

Lipids were analyzed on positive mode using a Bruker timsTOF fleX mass spectrometer coupled to a Bruker nanoElute 2 nanoLC system. Chromatographic separation was performed on a Bruker PepSep Ultra C18 column (25 cm × 75 μm, 1.5 μm particle size) using a 30 min gradient. Mobile phase A and B are 60% acetonitrile in water containing 10 mM ammonium formate, and isopropanol/acetonitrile (90:10, v/v) containing 10 mM ammonium formate, respectively. The flow rate was 250 nL/min, and the column oven temperature was maintained at 60 °C. Mass spectrometry data were acquired in PASEF mode over an m/z range of 100-1350. The ion mobility range was 1/K₀ = 0.55-1.79 V·s/cm², with a TIMS ramp time of 100 ms. The ion source was operated using the Bruker CaptiveSpray ion source with a capillary voltage of 1800 V and a source temperature of 180 °C. The collision energy and RF amplitude were set to 10 eV and 1100 Vpp, respectively. The quadrupole ion energy was 5 eV. The pre-TOF transfer time and pre-pulse storage time were 65 μs and 5 μs, respectively. Both m/z and ion mobility calibrations were performed prior to data acquisition using the Agilent ESI Low Concentration Tune Mix. The gas flow was adjusted to position the m/z 622 calibrant ion at 164 UV before ion mobility calibration. For post-acquisition mass recalibration, a calibration segment was acquired at the end of each run after gradient completion by injecting 2 μL of 1 mM sodium formate through the nanoLC system. All other instrument parameters were kept at default settings.

### LC-MS/MS data analysis

The LC-MSMS Raw data files were processed using Bruker Compass MetaboScape 2025 for feature finding and library-based MS/MS annotation. Feature extraction was performed using the MCube T-ReX 4D algorithm with an intensity threshold of 5,000. The [M+H]⁺ and [M+Na]⁺ ions were selected as candidate adducts for feature detection. Both mass recalibration and ion mobility recalibration were enabled during data processing. Lipid annotation was performed by searching against the NIST MSMS Spectral Library (https://chemdata.nist.gov/dokuwiki/doku.php?id=chemdata:start), the HMDB(*61*), and the MS-DIAL Tandem Mass Spectral Library for both positive and negative ionization modes(*62*). Library matching was restricted to MS/MS-confirmed only, using a 10 ppm mass tolerance and 5% ion mobility tolerance. All other parameters were maintained at default settings. Finally, annotated lipids meeting the criteria of ±10 ppm mass tolerance and ±5% CCS deviation were selected to generate a target list for subsequent MALDI-MSI data annotation.

### MALDI MSI data analysis

The MALDI-MSI data were analyzed using Bruker SCiLS Lab software (version 2025) for feature finding. The T-ReX algorithm with a 5 mDa bin size was applied for feature extraction. A 10 ppm mass interval width and a 0.05% intensity threshold were used during feature detection. The resulting .mca file was exported and imported into Bruker MetaboScape for target annotation using the kidney sample-specific target list derived from LC-MS/MS data generated using the same mass spectrometry platform, as described above. For MSI quantification, Root Mean Square normalization was applied. Final annotations were filtered using a ±10 ppm mass tolerance and ±5% CCS deviation for lipids successfully analyzed across compartments. All other parameters were kept at their default settings. Pixel-level MSI intensity data were exported using R scripts via the SCiLS Lab API function for further visualization and statistical analysis.

### Protein Sequence and Structure Acquisition

Human FBLN2 (UniProt ID: P98095) and EGFR (UniProt ID: P00533) sequences were retrieved from the UniProtKB database. FBLN2 is a 1184-amino acid extracellular matrix glycoprotein characterized by a central functional region containing 11 EGF-like domains. EGFR is a 1210-amino acid receptor tyrosine kinase; its extracellular domain (ECD, residues 1–645) contains Domains I and III, which form the canonical ligand-binding "sandwich". The crystal structure of the human EGFR complex was obtained from the RCSB Protein Data Bank (PDB ID: 7SYD). As no experimental structure or suitable homology template existed for FBLN2, its 3D structure was generated using AlphaFold3.

FBLN2 structural modeling was performed using a local implementation of the AlphaFold3 deep learning framework. Python and PyTorch-based dependencies were configured using the AlphaFold3 replication repository. Input tensors for pair and single representations were defined with a batch size of 1. Atomic coordinates were operated on using a diffusion module with 3 channels to predict protein folding. The AlphaFold3 model was initialized with a sequence length of 5, 8 attention heads, and 48 pair-former layers, running for 1000 diffusion steps to yield the final FBLN2 monomeric structure.

### Protein-Protein Docking

To explore the potential binding mode between the FBLN2 EGF-like domain and the EGFR ECD, molecular docking was performed using RosettaDock (Rosetta 3.13). The simulation followed a 1:2 stoichiometry, where one FBLN2 monomer bridges two EGFR subunits to induce receptor dimerization. Non-protein atoms were removed via clean_pdb.py, and missing side chains were repaired using fix_missing.xml. A 120 × 120 × 120 Å docking box was centered on FBLN2. A centroid-based Monte Carlo search (10,000 decoys) explored the interface using the cen_std scoring function. The top 1,000 models were clustered (radius 3.0 Å) to identify 200 candidate configurations. Candidates were converted to full-atom models using the Dunbrack Rotamer library. Local refinement involved small perturbations (±1.0 Å translation, ±3.0° rotation) and side-chain packing via PackRotamersMover. Interface scores (I_sc_ = E_bound_ - E_unbound_) were calculated using the talaris2014 energy function. The lowest-energy model (I_sc_= −15 kcal/mol) with a solvent-accessible surface area (SASA) > 800 Å² and > 3 interfacial hydrogen bonds was selected for MD simulation.

### Molecular Dynamics (MD) Simulation

All-atom MD simulations were performed using GROMACS 2020.4 with the AMBER ff99SB force field. The FBLN2/EGFR complex was placed in a 12 nm³ cubic box with TIP3P water. The system was neutralized with 13 Na⁺ ions to achieve a 0.15 M NaCl physiological concentration. A two-step steepest descent and conjugate gradient approach was used (5,000 steps each). Initial restraints (1,000 kJ/mol/nm²) on protein heavy atoms were applied and subsequently released to reach a maximum force < 100 kJ/mol/nm. The system underwent 100 ps of NVT ensemble heating to 300 K using Langevin dynamics, followed by 100 ps of NPT ensemble equilibration to 1 bar using the Berendsen barostat. A 200 ns production MD was conducted at 300 K and 1 bar using the Parrinello-Rahman barostat. Long-range electrostatics were treated via the Particle Mesh Ewald (PME) method. Trajectories were recorded every 10 ps. Structural stability was assessed by calculating the Root Mean Square Deviation (RMSD) relative to the initial equilibrated structure using the gmx rms tool.

### Gene expression analysis of FBLN2 or ACAT1 across human/mouse kidney cell types

To explore FBLN2 expression across kidney cell types, we searched for *FBLN2* in the KPMP Kidney Tissue Atlas (https://atlas.kpmp.org)(25). The expression of FBLN2 in healthy kidneys (N=40) and CKD kidneys (N=72) was plotted separately using single-cell neural RNA-seq data from KPMP. Cell type-specific expression of FBLN2 is determined by comparing it with all other cell types. FBLN2 and ACAT1 expression in mouse kidney cell types was mined from GSE139107(*24*).

### Analysis of regulatory variants and elements near the gene FBLN2

To explore the function of FBLN2 in kidney disease, we integrated the Genome-Wide Association Study (GWAS) of kidney function, measured by creatinine-based glomerular filtration rate, which included 2.2 million individuals(*27*). Significant variants associated with kidney function were defined as p < 5×10^-8^. The target genes of these variants were prioritized using the Kidney Disease Genetic Scorecard based on this GWAS (https://susztaklab.com/GWAS2M). The GWAS signals for the nearby genes FBLN2, FBLN1, and FBLN7 were visualized using LocusZoom(*63*). The chromatin accessibility around these genes was visualized using the Kidney Disease Genetic Scorecard.

### Expression changes of ACAT1 in AKI and CKD across human kidney cell types

The Human Kidney Single Cell Transcriptome by Abedini et al. was used to explore changes in ACAT1 expression in AKI and CKD across human kidney cell types(*30*). This atlas was built based on more than 200,000 cells from normal and diseased human kidneys. The differential expression between the normal and disease groups was assessed using scanpy’s rank_genes_groups function. P-values were calculated using a Wilcoxon rank-sum test and were corrected for multiple testing using the Benjamini-Hochberg method (adjusted for testing 34,733 genes).

### Primary fibroblasts isolation

Primary fibroblasts were isolated from the *Gli1-Smo*+/+ and *Gli1-Smo*-/- mice kidneys (10-day-old pups) and cultured with EMEM containing 10% FBS. *Smo* was deleted using (Z)-4-Hydroxytamoxifen (H7905, Sigma) for 5 days.

### Cell culture and treatment

NRK-49F (CRL-1570), and NRK-52E (CRL-1571) cells were obtained from ATCC. For conditioned media (CM) collections, under TGFβ stress, NRK-49F cells were incubated with CPN (2.5µM) or transfected with Dicer-substrate *Smo* siRNA (rn.Ri.Smo.13) for 24 h and then cultured with serum-free media for additional 24 h. The cultured medium was harvested and centrifuged (3000 rpm for 10 min at 4 °C). The supernatant was aliquoted and stored at −80°C for subsequent experiments. Serum-starved NRK-52E cells were then treated with the CM or Fibulin-2 (FBLN2) recombinant protein or transfected with Dicer-substrate Acat1 siRNA (rn.Ri.Acat1.13) under TGFβ stress.

### Fibroblast decellularized ECM scaffold preparation

Serum-starved NRK-49F were transfected with Dicer-substrate *Smo*-siRNA and treated with TGFβ for 24 hours. Cells were then treated with EGTA (#3889; Sigma) (0.5 mM, PH=7.4) in calcium-free PBS, followed by shaking at 4°C for 1 hour. The treatment was repeated 3-4 times until all cells were lifted from their underlying matrix. The fibroblast decellularized ECM scaffold was rinsed with PBS and then stored at 4°C for further experiments.

### Co-immunoprecipitation

NRK-52E cells were treated with rFBLN2 for 24 hours, cells lysates were incubated overnight at 4°C with 2 mg of anti-FBLN2 (sc-271843, Santa Cruz Biotechnology), followed by precipitation with 100 µl of protein A/G Plus-agarose for 3 hours at 4°C. The precipitated complexes were separated by SDS–polyacrylamide gel electrophoresis and immunoblotted with specific antibodies EGFR (18986-1-AP, Proteintech Group), respectively.

### Automated quantitation for immunostaining

For each kidney section, five images per mouse (five mice/group) were randomly selected for analysis. Positive staining area was quantified using a custom script in Image Pro plus 6.0(*64*). Images were separated into red, green, and blue channels, and the channel with the best signal-to-background separation was used for analysis. Thresholding (0, 80) was manually optimized to highlight high-intensity marker expression and applied uniformly across all images for a given marker. The percentage of threshold-positive area was measured and used as an index of marker expression.

### Statistics

Statistical analyses were performed using GraphPad Prism 9. Data distribution was assumed to be normal but this was not formally tested. Comparisons between two groups used a two-sided Student’s t-test or a two-sided Wilcoxon test, while multiple-group comparisons were analyzed by one-way ANOVA followed by the Student-Newman-Keuls test. Results are presented in dot plots, with each dot denoting an individual value. *P* < 0.05 was considered statistically significant. For lipidomics, Differential lipid analysis was performed using the limma package(*65*). Lipids were considered significantly different if they met the thresholds of fold change better than 1.2 and adjusted *P* value < 0.05.

## Data Availability

All unique materials generated in this study are available from the corresponding author upon completion of a Materials Transfer Agreement. Raw mass spectrometry data have been deposited in the ProteomeXchange Consortium via the PRIDE partner repository with the dataset identifier PXD077785.

## Author contributions

Conceptualization: GY, ZD; Methodology: GY, WYY, LWX, LS, LHB, YYB, CW; Investigation: GY, WYY, LWX, DC, LJJ, MS, ZK, JC, SWH, DYL, CT, MB; Visualization: GY, LWX, ZD; Funding acquisition: LYS, ZD; Project administration: ZD; Supervision: ZD; Writing – original draft: GY, ZD; Writing – review & editing: GY, LYS, ZD. The co-first authorship order reflects contribution to the study.

## Acknowledgments

We are grateful to Dr. Anna Mae Diehl at Duke University for generously providing the *Smo*-flox mice. We also thank Kidney Precision Medicine Project, Dr. Benjamin Humphreys’s laboratory at Washington University in St. Louis, and Dr. Katalin Susztak’s laboratory at University of Pennsylvania for making their publicly available datasets accessible for this study. We thank Bernard L. Cook, a science editor and illustrator at UConn Health, for creating artwork for this manuscript.

