## Supplementary Figures and Table 2-3 for "Fibulin-2 transduces a matrix-to-metabolism signal in kidney fibrosis"

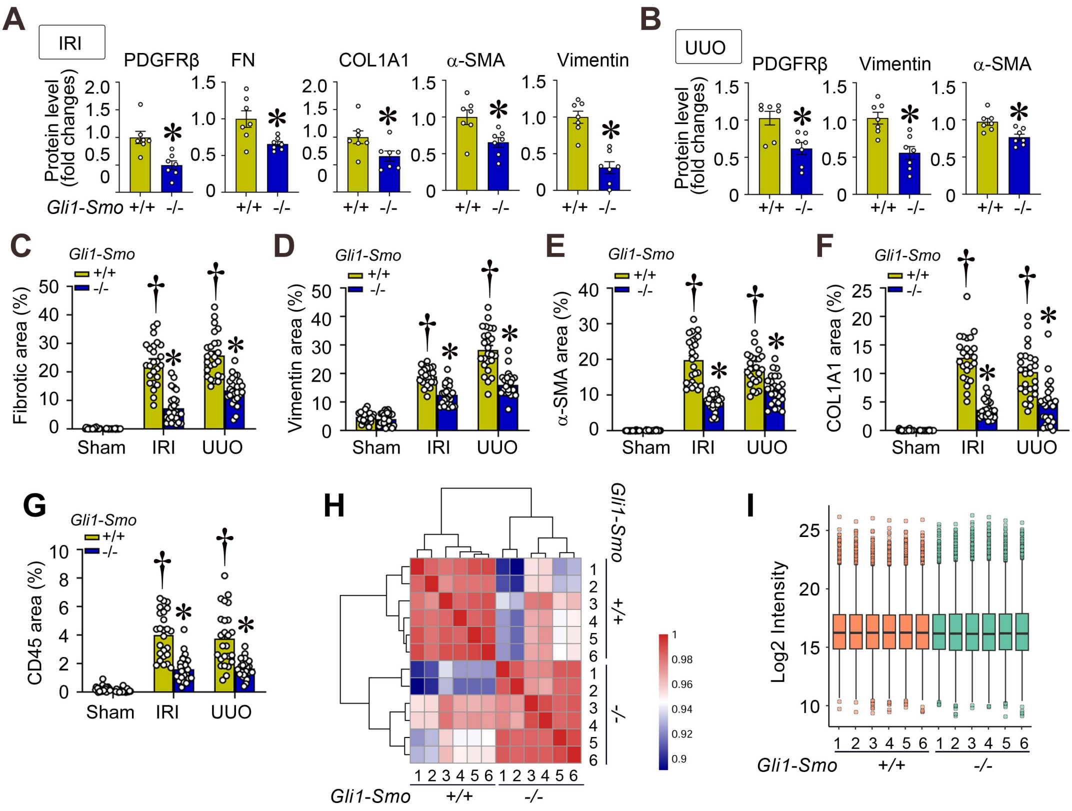

**Supplementary Figure S1: Global proteomics reveals FBLN2 regulates kidney metabolism after loss of *Smo* in fibroblasts in fibrotic kidneys.** **(A)** Quantitative data for PDGFRβ, FN, COL1A1, and α-SMA after unilateral IRI combined with nephrectomy (n = 7). (**B**) Quantitative data for PDGFRβ, Vimentin, and α-SMA after unilateral ureteral obstruction (n = 7). (**C-G**) Quantitative data for MTS (**C**) and immunohistochemical staining for Vimentin (**D**), α-SMA (**E**), COL1A1 (**F**), and CD45 (**G**) in ischemic or obstructive kidneys between *Gli1-Smo*+/+ and *Gli1-Smo*-/- kidneys (n = 5. 5 random images were selected per mouse; each dot represents the score of the according image). (**H**) Replicate correlation of the kidney proteome profiles between *Gli1-Smo*+/+ and *Gli1-Smo*-/- kidneys after unilateral IRI combined with nephrectomy. (**I**) Violin plot of significant proteins (Permutation FDR 0.05) among the *Gli1-Smo*+/+ and *Gli1-Smo*-/- mice. LFQ intensity of represented proteins was z-scored and plotted according to the color bar.† P < 0.05 versus sham control, * P < 0.05 versus *Gli1-Smo*+/+ mice after UIRI+UNx or UUO. Graphs are presented as means ± SEM. Differences among groups were analyzed using two-sided unpaired t-tests or one-way ANOVA followed by the Student-Newman-Keuls test.

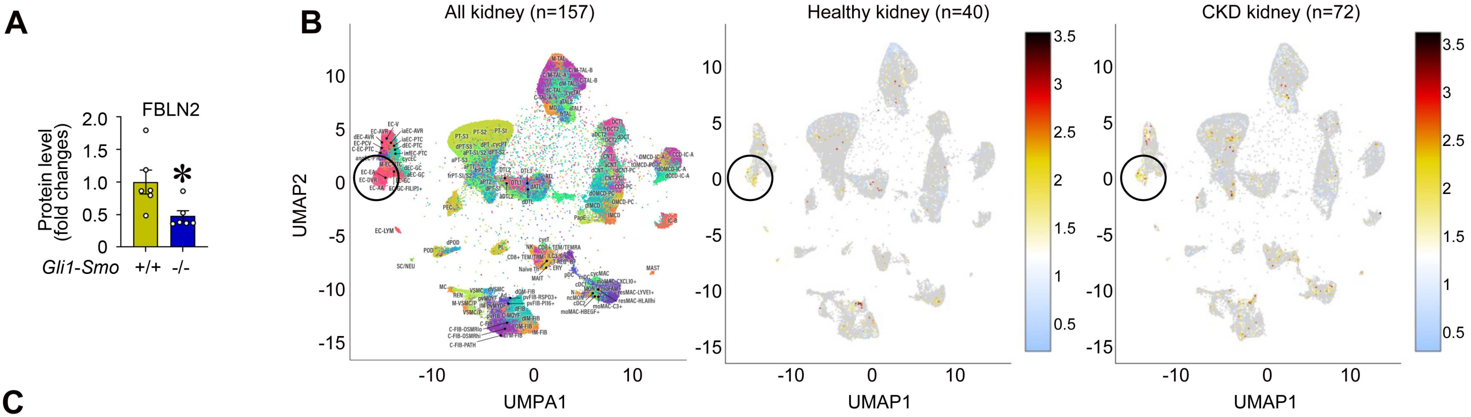


C

| clusterAbbrev | clusterName | cellCount | meanExp | pctCellsExpressing | foldChange | pVal | pValAdj |
|---|---|---|---|---|---|---|---|
| EC-AA | Afferent Artery / Arteriole Endothelial Cell | 1440 | 2.58 | 43.5 | 6.02 | 1.86E-214 | 6.80E-210 |
| EC-DVR | Descending Vasa Recta Endothelial Cell | 4598 | 1.92 | 36.2 | 5.92 | 5.42E-186 | 1.98E-181 |
| pvFIB-PI16+ | Perivascular Fibroblast, PI16+CD34+ | 977 | 1.44 | 30.2 | 5.05 | 7.04E-131 | 2.57E-126 |
| Ad | Adipocyte | 129 | 0.693 | 19.4 | 3.94 | 8.13E-37 | 2.97E-32 |
| SC/NEU | Schwann Cell / Neural | 229 | 0.629 | 10.5 | 3.8 | 9.43E-21 | 3.45E-16 |

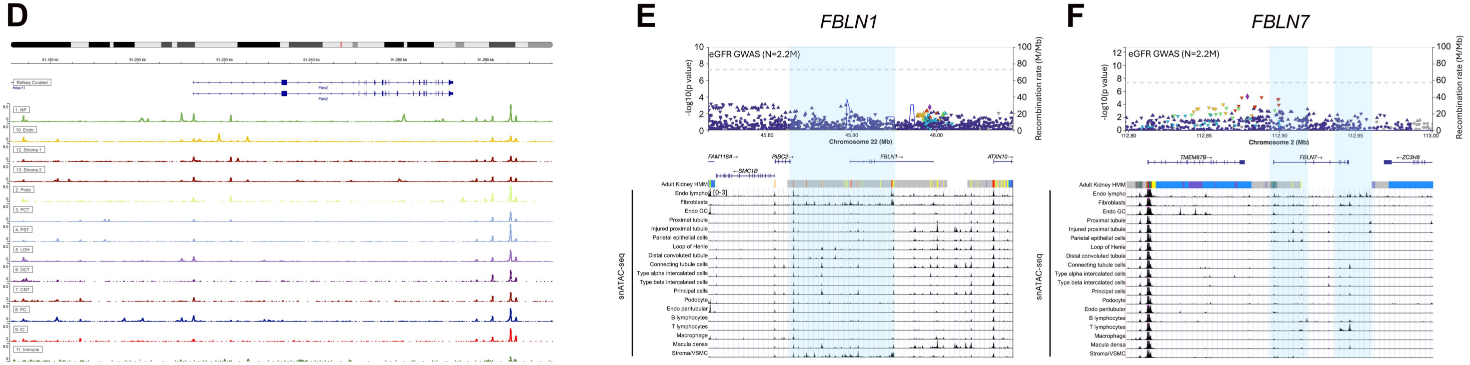


**Supplementary Figure S2: FBLN2 is a key core matrisome component expressed in fibroblasts and endothelial cells. (A)** Quantitative data of FBLN2 expression in the fibrotic kidneys between *Gli1-Smo*+/+ and *Gli1-Smo*-/- mice (n = 6). (**B, C**) KPMP datasets (merged with datasets from Dr. Susztak Katalin's laboratory) showing fibroblasts and endothelial cells are major cellular sources of FBLN2 in kidneys from CKD patients. (**D**) Fbln2 promoter showing higher open chromatin in fibroblasts and endothelial cells in the rat kidney. (**E, F**) Genetic ScoreCard of GWAS variants in the gene FBLN1 (**E**) and FBLN7 (**F**) locus. The panels from top to bottom show the eGFR GWAS mapped in 2.2 million individuals, genes, variant-to-gene associations based on methylation QTL, chromatin state in adult kidney, and open chromatin in kidney cell types profiled by snATAC-seq. * $P < 0.05$. Graphs are presented as means ± SEM. Differences among groups were analyzed using two-sided unpaired t-tests.

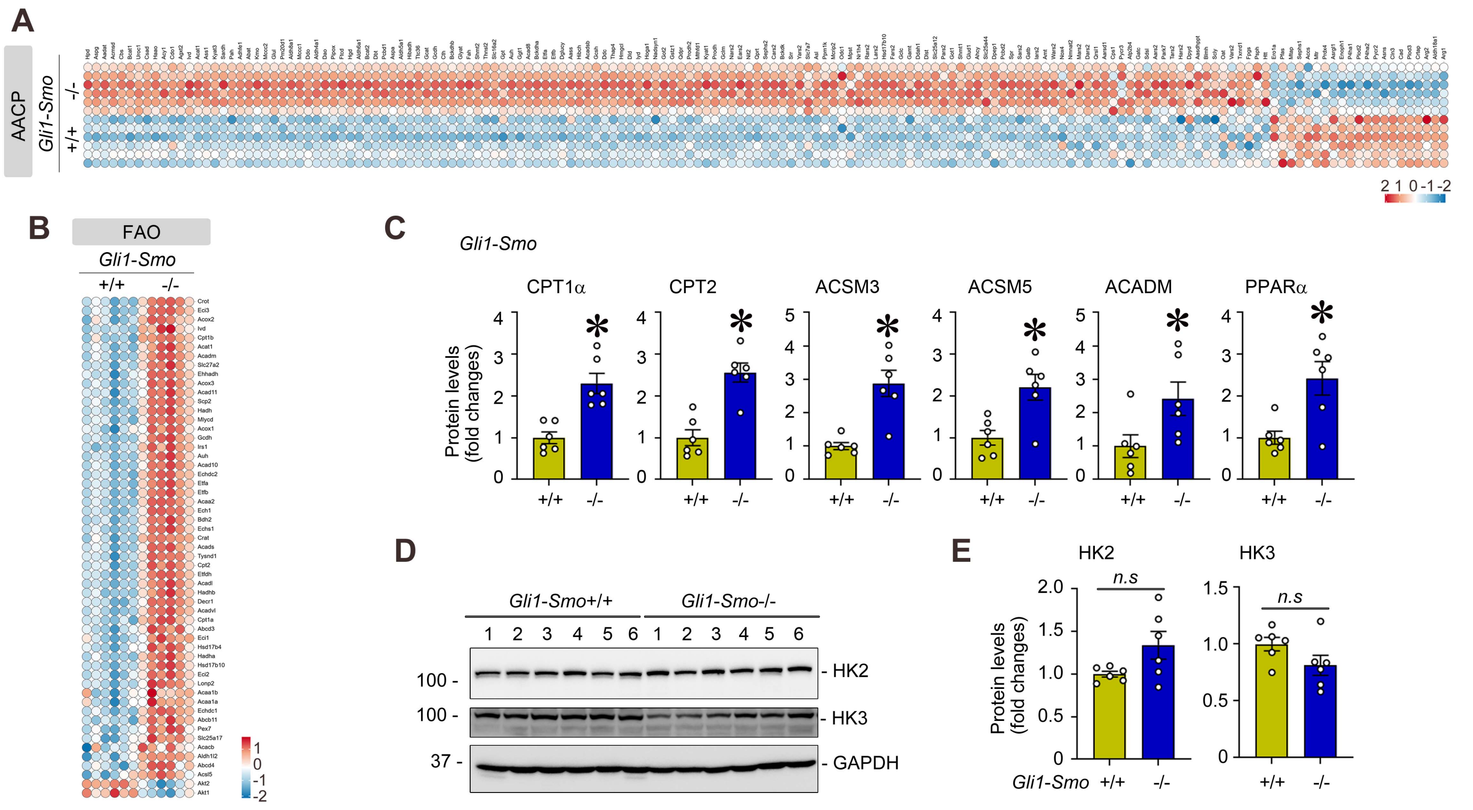


**Supplementary Figure S3: Fibroblast-specific deletion of *Smo* enhances fatty acid oxidation and amino acid metabolism in fibrotic kidneys. (A, B)** Heatmap of the differentially expressed proteins in the amino acid metabolism (**A**), and Fatty acid oxidation (FAO) (**B**) between *Gli1-Smo*+/+ and *Gli1-Smo*-/- kidneys after unilateral IRI combined with nephrectomy. (**C**) Quantitative data of CPT1α, CPT2, ACSM3, ACSM5, ACADM, and PPARα proteins in fibrotic kidneys from both group (n = 6). (**D, E**) Western blot (**D**) and quantitative data (**E**) of HK2 and HK3 proteins (n = 6). * $P < 0.05$. Graphs are presented as means ± SEM. Differences among groups were analyzed using two-sided unpaired t-tests.

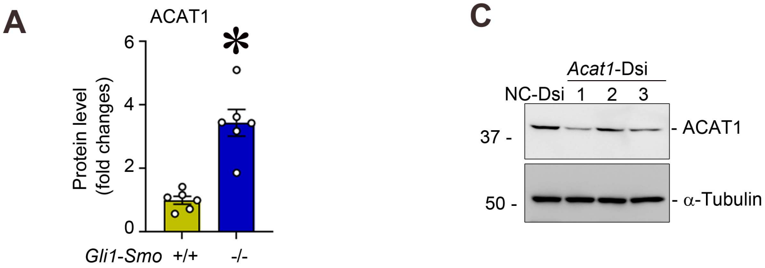


**B**

| Cell type | Average Expression (All Samples) | Fraction Expressing (All Samples) | CellType marker p-values | Average Expression (Control Samples) | Average Expression (AKI Samples) | Average Expression (CKD Samples) | Fraction Expressing (Control Samples) | Fraction Expressing (AKI Samples) | Fraction Expressing (CKD Samples) | AKI vs. Control p-values | CKD vs. Control p-values |
|---|---|---|---|---|---|---|---|---|---|---|---|
| PT_S2 | 1.33 | 0.59 | 0.00E+00 | 1.45 | 1.07 | 1.37 | 0.62 | 0.53 | 0.58 | 3.40E-150 | 6.40E-39 |
| PT_S1 | 1.28 | 0.62 | 0.00E+00 | 1.32 | 1.3 | 1.25 | 0.65 | 0.67 | 0.58 | 8.00E-07 | 4.50E-09 |
| PT_S3 | 1.28 | 0.61 | 0.00E+00 | 1.35 | 1.15 | 1.33 | 0.62 | 0.63 | 0.61 | 9.60E-26 | 1.00E-05 |
| IC_A | 1.2 | 0.72 | 0.00E+00 | 1.17 | 1 | 1.26 | 0.73 | 0.64 | 0.7 | 5.40E-06 | 4.90E-34 |
| C_TAL | 0.72 | 0.47 | 0.00E+00 | 0.77 | 0.65 | 0.7 | 0.54 | 0.42 | 0.42 | 9.50E-04 | 2.50E-05 |

**Supplementary Figure S4: ACAT1 is a common regulatory node mediating fatty acid oxidation and amino acid metabolism in fibrotic kidneys. (A)** Quantitative data of ACAT1 protein in *Gli1-Smo*+/+ and *Gli-Smo*-/- kidneys after unilateral IRI combined with nephrectomy (n = 6). (**B**) KPMP datasets (merged with datasets from Dr. Susztak Katalin's laboratory) showing tubular cells are major cellular sources of ACAT1 in kidneys from CKD patients. (**C**) Western blot analyses confirm the efficiency of ACAT1 knockdown in NRK-52E cells using three individuals predesigned DsiRNA (1, 2, and 3). NC, negative control siRNA. * $P < 0.05$. Graphs are presented as means ± SEM. Differences among groups were analyzed using two-sided unpaired t-tests.

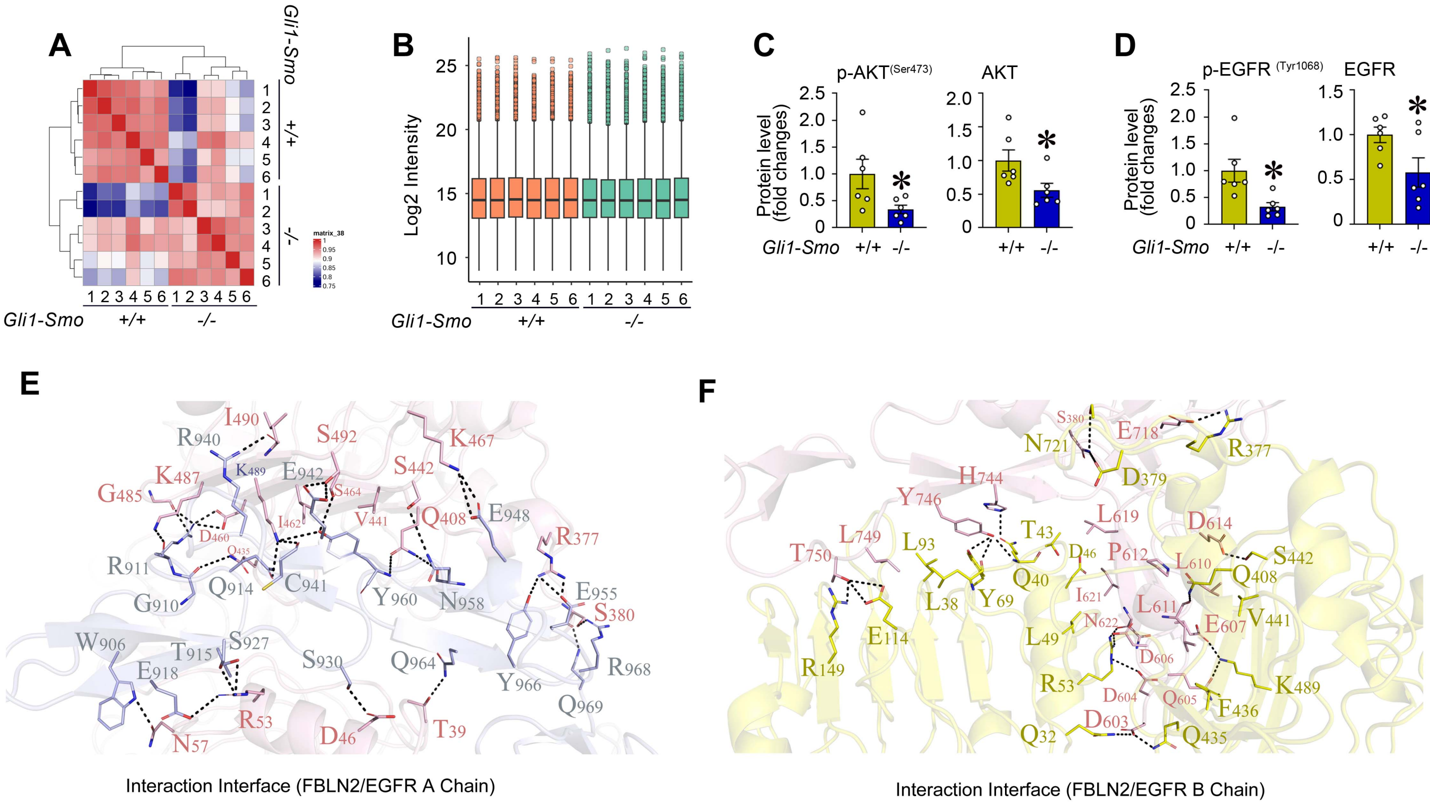


**Supplementary Figure S5: Phosphoproteomics reveals suppressed EGFR/AKT in tubules after Smo deletion in fibroblasts. (A)** Replicate correlation and the density distributions of the kidney phosphoproteome profiles between *Gli1-Smo*+/+ and *Gli1-Smo*-/- kidneys after unilateral IRI combined with nephrectomy. (**B**) Violin plot of significant proteins (Permutation FDR 0.05) among the *Gli1-Smo*+/+ and *Gli1-Smo*-/- mice. LFQ intensity of represented proteins was z-scored and plotted according to the color bar. (**c**, **d**) Quantitative data of p-AKT (Ser473), AKT (n = 6) (**C**), and p-EGFR (Tyr1068), EGFR proteins (n = 6) (**D**) in *Gli1-Smo*+/+ and *Gli1-Smo*-/- kidneys after unilateral IRI combined with nephrectomy. (**E, F**) Protein-protein docking showing the interaction interface of FBLN2 and EGFR Chain A (**E**) and Chain B (**F**). * P < 0.05. Graphs are presented as means ± SEM. Differences among groups were analyzed using two-sided unpaired t-tests.

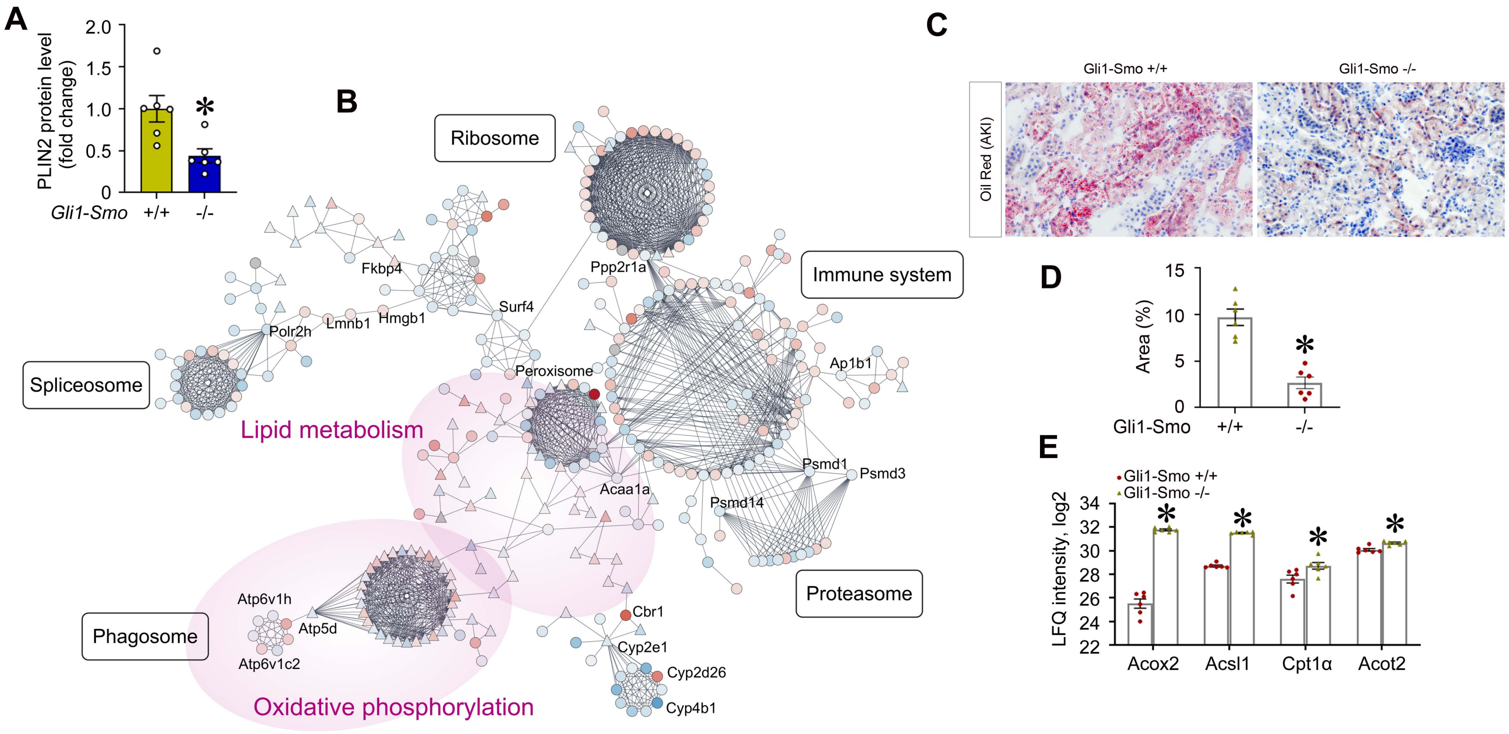


**Supplementary Figure S6: Fibroblast-specific deletion of *Smo* impacts fatty acid oxidation after AKI. (A)** Quantitative data of PLIN2 protein in *Gli1-Smo*+/+ and *Gli1-Smo*-/- kidneys after UIRI+UNx. (n = 6). **(B)** Lipid associated network in *Gli1-Smo*+/+ and *Gli1-Smo*-/- kidneys after AKI. The network and enrichment results were derived from String App in Cytoscape software. (**C**) Oil-Red staining in *Gli1-Smo*+/+ and Gli1-Smo-/- kidneys after acute kidney injury (AKI). (**D**) Quantitative data for Oil-Red staining. (**E**) Protein intensity in fatty acid oxidation pathway from our global proteomics data of AKI. * $P < 0.05$. Graphs are presented as means ± SEM. Differences among groups were analyzed using two-sided unpaired t-tests.

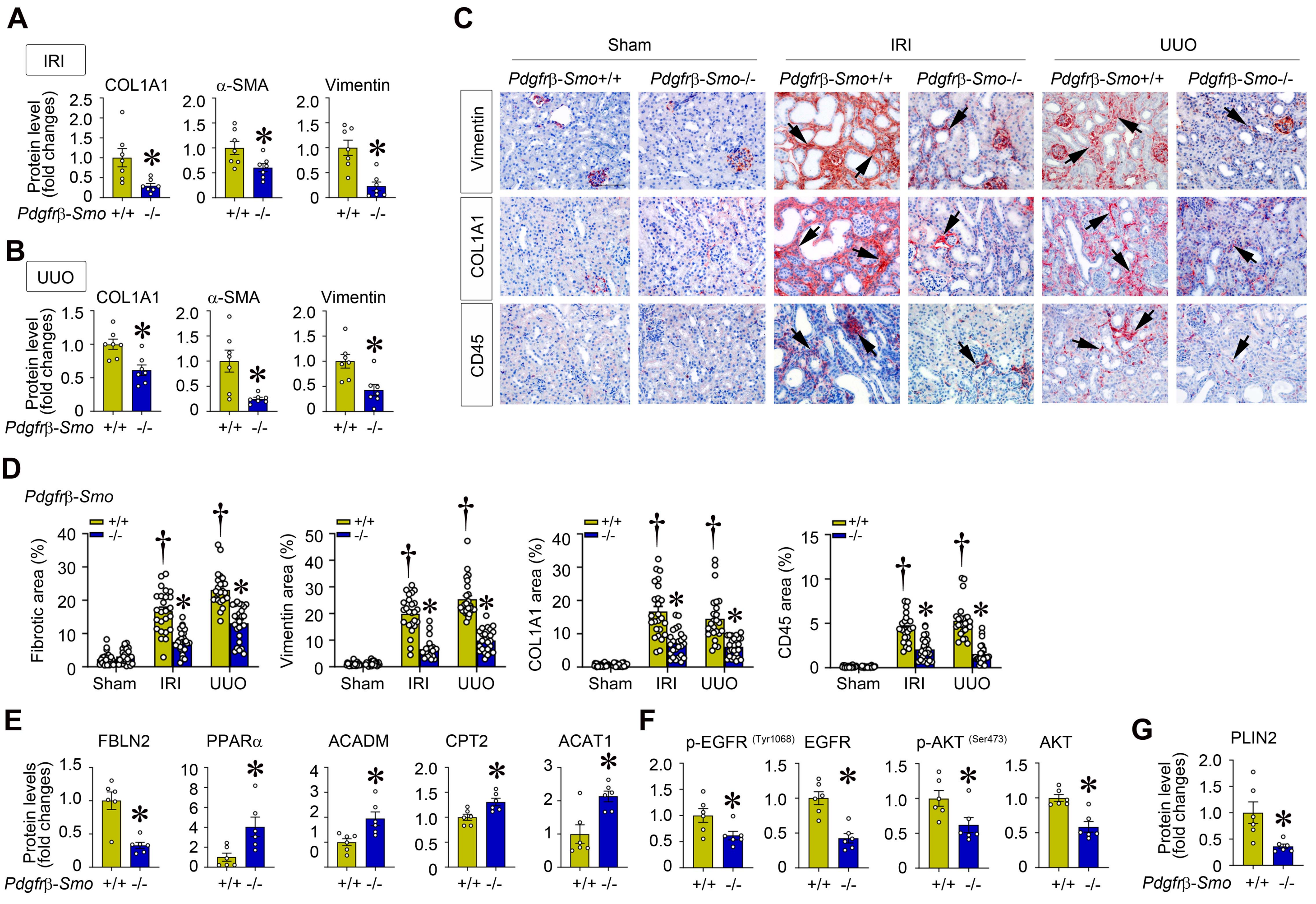


**Supplementary Figure S7: Loss of *Smo* in *Pdgfrb*+ fibroblasts mitigates kidney fibrosis after IRI and UUO.** In *Pdgfrβ-Smo*+/+ and *Pdgfrβ-Smo*-/- kidneys, after UIRI combined with nephrectomy or UUO, **(A, B)** quantitative data of COL1A1, α-SMA, and Vimentin proteins (n = 7), **(C)** Immunohistochemical staining for Vimentin, COL1A1, and CD45, **(D)** quantitative data of Masson's Trichrome staining (MTS) and immunohistochemistry (Vimentin, COL1A1, CD45) (n = 5. 5 random images were selected per mouse; each dot represents the score of the according image). In *Pdgfrβ-Smo*+/+ and *Pdgfrβ-Smo*-/- kidneys, after UIRI combined with nephrectomy, **(E-G)** quantitative data of FBLN2, PPARα, ACADM, CPT2, ACAT1 (n = 6) **(E)**, p-EGFR (Tyr1068), EGFR, p-AKT (Ser473), AKT (n = 6) **(F)**, PLIN2 (n = 6) **(G)** proteins. † P < 0.05 versus sham control, * P < 0.05 versus Pdgfrβ-Smo+/+ mice after UIRI+UNx or UUO. Graphs are presented as means ± SEM. Differences among groups were analyzed using two-sided unpaired t-tests or one-way ANOVA followed by the Student-Newman-Keuls test.

**Supplementary Table S2. The information of the applied primary and secondary antibodies**

| Name | Vendor | Category Number | Application |
|---|---|---|---|
| PDGFR-β | Cell signaling Technology, Danvers, MA | #3169 | WB |
| Fibronectin | Sigma, St. Louis, MO | F3648 | WB |
| COL1A1 | Cell signaling Technology, Danvers, MA | #72026 | WB, IHC |
| α-SMA | Abcam, Cambridge, MA | ab5694 | WB, IHC |
| Vimentin | Cell signaling Technology, Danvers, MA | #5741 | WB, IHC |
| CD45 | Cell signaling Technology, Danvers, MA | #70257 | IHC |
| FBLN2 | Santa Cruz Biotechnology, Dallas, Texas | sc-271843 | WB, CO-IP |
| CPT1α | Proteintech Group, Rosemont, IL | 15184-1-AP | WB, IHC |
| CPT2 | Proteintech Group, Rosemont, IL | 26555-1-AP | WB |
| ACSM3 | Proteintech Group, Rosemont, IL | 10168-1-AP | WB |
| ACSM5 | Proteintech Group, Rosemont, IL | 16591-1-AP | WB |
| ACADM | Proteintech Group, Rosemont, IL | 55210-1-AP | WB |
| PPARα | Proteintech Group, Rosemont, IL | 15540-1-AP | WB, IHC |
| ACAT1 | Proteintech Group, Rosemont, IL | 16215-1-AP | WB, IHC |
| P-ACAT1 (Try407) | Signalway Antibody LLC (SAB), Greenbelt, Maryland, | SAB491P | WB |
| p-EGFR (Try1068) | Santa Cruz Biotechnology, Dallas, Texas | sc-81488 | WB |
| EGFR | Santa Cruz Biotechnology, Dallas, Texas | sc-373746 | WB |
| p-EGFR(Try1068) | Cell signaling Technology, Danvers, MA | #3777 | WB |
| EGFR | Cell signaling Technology, Danvers, MA | #4267 | WB |
| EGFR | Proteintech Group, Rosemont, IL | 18986-1-AP | WB |
| PLIN2 | Abcam, Cambridge, MA | ab52356 | WB, IF |
| p-AKT (Ser473) | Cell signaling Technology, Danvers, MA | #4060 | WB |
| AKT | Cell signaling Technology, Danvers, MA | #4691 | WB |
| HK2 | Invitrogen, Thermo Fisher Scientific, MA | PA5-293626 | WB |
| HK3 | Invitrogen, Thermo Fisher Scientific, MA | PA5-76388 | WB |
| α-Tubulin | Sigma, St. Louis, MO | T9026 | WB |
| GAPDH | Santa Cruz Biotechnology, Dallas, Texas | sc-32233 | WB |
| Anti-Mouse IgG | Abcam, Cambridge, MA | ab6789 | WB |
| Anti-Rabbit IgG | Abcam, Cambridge, MA | ab6721 | WB |
| Alexa Fluor® 488 Anti-Rabbit | Jackson ImmunoResearch | 711-545-152 | IF |
| Biotin-Anti-Rabbit | Jackson ImmunoResearch | 711-065-152 | IHC |

WB, western blot; CO-IP, Co-Immunoprecipitation; IHC, Immunohistochemical staining; IF, Immunofluorescence staining

**Supplementary Table S3. Nucleotide sequences of the primers used for qRT-PCR**

| Mouse gene | Primer Sequence 5' to 3' | |
|---|---|---|
| | Forward | Reverse |
| IL-6 (M) | CTTGGGACTGATGCTGGTG | TCCACGATTTCCCAGAGAAC |
| MCP1 (M) | TTAAAAACCTGGATCGGAACCAA | GCATTAGCTTCAGATTTACGGGT |
| TNFα (M) | CCCTCACACTCAGATCATCTTCT | CCCTCACACTCAGATCATCTTCT |
| Cd36 (M) | GGACATTGAGATTCTTTTCCTCTG | GCAAAGGCATTGGCTGGAAGAAC |
| Cpt2 (M) | GATGGCTGAGTGCTCCAAATACC | GCTGCCAGATACCGTAGAGCAA |
| Acadl (M) | GGCGATTTCTGCCTGTGAGTTC | GCTGTCCACAAAAGCTCTGGTG |
| ACOX3 (M) | CCTATGCCTTGGACCACTTCTC | ATGCCAGAGCATGGATCTCACG |
| Acsm5 (M) | TCTTCTCTGCCTGGTCCAATGG | AAGAGGGTTGGGACACAGCACA |
| Acadm (M) | AGGATGACGGAGCAGCCAATGA | GCCGTTGATAACATACTCGTCAC |
| Glul (M) | CTGCCATACCAACTTCAGCACC | CTGGTGCCTCTTGCTCAGTTTG |
| Gucy2d (M) | GAGAACCTGAGGCTAGACTGGA | AGCACAAAGCGAGTGTCCACCA |
| Glud1(M) | TCCGTTACAGCACTGACGTGAG | ACGCCTGCTTTAGCACCTCCAA |
| Slc25A21 (M) | CGAGGTGGTAAAAGTTGGCTTGC | GCTGTCAATCCTTTGTCGAGGC |
| Slc25A44 (M) | CTCGCTGCTAACGTACATCCCA | GGCTTGAAAGACAATGTGAGGGC |
| Ppara (M) | ACCACTACGGAGTTCACGCATG | GAATCTTGCAGCTCCGATCACAC |
| Acox1 (M) | GCCATTCGATACAGTGCTGTGAG | CCGAGAAAGTGGAAGGCATAGG |
| Acat1 (M) | GCAGGGAAGTTTGCCAGTGAGA | GAACACGGTCTTGAGCTTTGGC |
| Col1α1(M) | ATCTCCTGGTGCTGATGGAC | ACCTTGTTTGCCAGGTTCAC |
| Col3α1 (M) | AGGCAACAGTGGTTCTCCTG | GACCTCGTGCTCCAGTTAGC |
| Fbln2 (M) | AGCCAGGCTATGTCCTCACAGA | GTAGAAGGAGCCCTTGGTGTTC |
| Plin2 (M) | GACAGGATGGAGGAAAGACTGC | GGTAGTCGTCACCACATCCTTC |
| Glud1(Rat) | AGATTCACCATGGAGCTGGC | ATGGTGCTGGCATAGGTGTC |
| Slc25A44 (Rat) | CAAGTGCGTGGGAACCTAGA | GCTGCTCTGCATAGAAGTGGTA |
| Cpt2 (Rat) | CTAAGAGATGCTCCGAGGCG | CAAGTGTCGGTCAAAGCCCT |
| Acox1 (Rat) | CTCACTCGAAGCCAGCGTTA | TTGAGGCCAACAGGTTCCAC |
| PPARα (Rat) | ACGATGCTGTCCTCCTTGATG | GCGTCTGACTCGGTCTTCTTG |
| Acadl (Rat) | AGTGCCCTACTTGGGGAAGA | CACAGTCTGGATGTGTGCGA |
| Acat1 (Rat) | ACCCGAAGTAAAGAGGCGTG | GAGCTCCAGACATCCCGATT |
| β-actin (M) | CAGCTGAGAGGGAAATCGTG | CGTTGCCAATAGTGATGACC |
| β-actin (Rat) | TTCCTGGGTATGGAATCCTG | CTTCTGCATCCTGTCAGCAA |
